# *TAOK3* inhibition constrains invasion, potentiates paclitaxel, and reprograms the tumor microenvironment toward anti-tumor immunity in cervical cancer

**DOI:** 10.64898/2026.07.04.736128

**Authors:** Marissa Iden, Rachel Schmidt, Rameesa Darul Amne Syed Mohammed, Theresa A. Dlugi, Roshan Kumar, Shirng-Wern Tsaih, Bakhtiyor Nosirov, Ishaque P. Kadamberi, Sonam Mittal, Shruti L. Narayan, William H. Bradley, Beth Erickson, Rebecca C. Czaja, Juan C. Felix, Victor Jin, Akinyemi I. Ojesina, Sunila Pradeep, Brian C. Smith, Janet S. Rader

## Abstract

*TAOK3* is a lesser-studied MAPK family serine/threonine kinase our group has shown to be targeted by HPV integration, suggesting a potential role in driving invasive cervical cancer (ICC). Here, we profiled *TAOK3* expression in patient tumors, metastases, and cervical cancer models and localized *TAOK3* within a tumor epithelial subpopulation by integrating two single-cell RNA-seq datasets. Functional consequences of *TAOK3* loss were assessed with siRNA and CRISPRi in cell lines and 3D spheroids. *In vivo* effects were evaluated in intracervical xenografts with species-specific RNA-seq to resolve tumor versus microenvironmental responses. *TAOK3* mRNA/protein were elevated in primary and metastatic ICC and primarily localized to a keratin-positive epithelial subset (T3epi) enriched for cadherin/S100 binding, vesicle/endocytic pathways, and leading-edge programs. *TAOK3* silencing reprogrammed transcriptomes and proteomes toward reduced WNT/cell-cycle and motility signaling, altered endocytosis and cytoskeleton organization, and reshaped phospho-networks linked to chromatin remodeling and ERBB2–ERBB3/cytoskeletal kinase activity. Functionally, *TAOK3* inhibition prolonged G2/M, suppressed invasion, and enhanced sensitivity to low dose paclitaxel. Prolonged inactivation induced methuosis-like cell death with extracellular ATP release. In xenografts, *TAOK3* knockdown reduced tumor burden, downregulated *KRT14*—a leader cell marker—within the human tumor compartment, and enriched microenvironmental pathways for immune activation, with a specific decrease in CD206+ M2 macrophages. *TAOK3* delineates an invasion-competent epithelial state in ICC and coordinates cell-cycle control, cytoskeleton–membrane dynamics, and tumor–immune crosstalk. Genetic or pharmacologic *TAOK3* inhibition constrains tumor growth, potentiates paclitaxel, and remodels the microenvironment toward anti-tumor immunity, supporting *TAOK3* as a potential therapeutic target and biomarker in ICC.

**Statement of Significance:** *TAOK3* marks an invasion-competent epithelial subpopulation in cervical cancer. *TAOK3* inhibition slows tumor growth, enhances chemoresponse, and reduces M2 macrophages, revealing *TAOK3* as a potential therapeutic target and biomarker for patient stratification.

## Introduction

Cervical cancer remains a leading cause of cancer-related morbidity and mortality worldwide, with persistent infection by high-risk human papillomaviruses (HPV) as the principal etiologic driver. Despite advances in prophylactic vaccination and screening, many women present with or progress to invasive cervical cancer (ICC), where limited therapeutic options and resistance to standard regimens constrain outcomes. First-line therapy for recurrent or metastatic disease typically combines platinum agents, paclitaxel, and sometimes bevacizumab and immune checkpoint blockade; however, durable responses remain uncommon, and treatment-limiting toxicities are frequent. These challenges underscore the need to identify tumor-intrinsic programs that enable invasion, immune evasion, and therapy resistance, and to develop biomarkers and targets that can be leveraged to improve clinical benefit.

HPV integration into the human genome is a critical process driving cervical cancer progression (1,2). HPV integration most often occurs in open, easily accessible regions of the host genome, suggesting that these regions likely harbor genes involved in cervical carcinogenesis. Indeed, recurrent HPV integrations have been identified in regions containing known oncogenes, such as *MYC/PVT1* (3) and TP63 (4). Even so, HPV integration at well-studied oncogenes accounts for less than 10% of reported integration events. Thus, regions of HPV integration may help illuminate potential, therapeutically relevant cervical cancer target genes. Our laboratory recently reported potential therapeutic target genes for cervical cancer based on long-read sequencing of HPV integration events (5). One of these genes, thousand and one amino acid kinase 3 (*TAOK3*), is a serine/threonine protein kinase and member of the germinal center kinase (GCK) subfamily of STE20-like kinases, which participate in multiple pathways regulating cell proliferation, apoptosis, actin-based cell protrusions like invadopodia, and immune response (6). Adding to its potential importance in promoting virus-mediated carcinogenesis, Taok3 was 1 of 19 loci found to be associated with mouse hepatocellular carcinoma based on a transposon-based insertional mutagenesis screen (7).

Mitogen-activated protein kinase (MAPK) signaling integrates extracellular stress and growth cues to regulate proliferation, survival, cytoskeletal dynamics, and differentiation, all of which are central to epithelial tumor progression. The role of *TAOK3* in cancer appears complex and potentially tissue- and/or cell-state-specific. For example, *TAOK3* can activate p38 and/or ERK (8,9) and can either inhibit or promote JNK activation in response to epidermal growth factor (EGF;(10)) or endoplasmic reticulum (ER) stress (11), respectively. *TAOK3* is also highly expressed in immune cells, particularly macrophages, T-cells, and dendritic and natural killer cells (6). Here, we combine normal cervix and tumor tissue profiling, integrative single-cell analyses, and multi-omic perturbations to uncover a TAOK3-high, keratin-positive epithelial subpopulation (T3epi) enriched for cadherin/S100 binding and vesicle/endocytic programs consistent with a motile, invasion-competent state. Through transcriptomic, proteomic, and phosphoproteomic profiling, along with functional assays, we show that *TAOK3* coordinates cell-cycle progression, cytoskeleton–membrane dynamics, and tumor–immune interactions, including chemosensitivity to paclitaxel and macrophage polarization *in vivo*. These findings position *TAOK3* as a mechanistic node linking leader cell behavior and microenvironmental remodeling in ICC and motivate therapeutic strategies that target *TAOK3* to constrain malignant progression and enhance current chemotherapy.

## Materials and Methods

### Cell culture

Four ICC cell lines, 1 commercially available (SiHa) and 3 patient-derived, primary cell lines (MCW-1, MCW-2, and MCW-3) were used in this study. SiHa (ATCC; HTB-35) were grown in high glucose Dulbecco’s Modified Eagle’s Medium (DMEM; ThermoFisher) supplemented with 10% v/v Fetal Bovine Serum (FBS). MCW-1, MCW-2, and MCW-3 patient-derived ICC cell lines were generated by our laboratory (see (12) and supplementary methods) and grown in F-medium (3:1 v/v F-12 [Ham]-DMEM; Invitrogen, ThermoFisher), 2.5% FBS, 0.4 μg/mL hydrocortisone (Millipore Sigma), 5 μg/mL insulin (Millipore Sigma), 8.4 ng/mL cholera toxin (Millipore Sigma), 10 ng/mL EGF (R&D Systems), 24 μg/mL adenine (Millipore Sigma), 100 U/mL penicillin, and 100 μg/mL streptomycin. For 3D cultures, 5,000 cells/well were seeded in a 96-well Nunclon Sphera, U-bottom plate (ThermoFisher) and centrifuged at 70 × g for 5 min before placing in the incubator. Media replacement was performed every 2–3 days, and spheroid formation was monitored for 7–9 days. Organoid cultures were grown according to a previously published protocol (13). All cultures were kept at 37°C in a high-humidity environment with a gas mixture consisting of 95% air and 5% CO_2_. Cell line authenticity was verified through STR profiling (ATCC), and cells were regularly checked for *Mycoplasma* using the PCR-based Universal *Mycoplasma* Detection Kit (ATCC) or the MycoStrip™ detection kit (InvivoGen). Cell harvesting was performed using a 0.25% Trypsin-EDTA solution and cell counting carried out using the Countess-3 Automated Cell Counter (Invitrogen) according to the manufacturer’s protocol. The study protocol for use of patient samples was approved by MCW’s Institutional Review Board.

### Transfections

SiHa, MCW-1, MCW-2, or MCW-3 cells were seeded (2.0⍰ × ⍰10^5^ cells/ml) into a 6-well plate (1 ml) or 10 cm dish (5 ml) and allowed to adhere overnight. Cells were transfected the next day with Qiagen AllStars Negative Control siRNA (siCONT) or Hs_TAOK3_4 FlexiTube (siTAOK3) siRNAs using DharmaFECT1 (Horizon Discovery, Waterbeach, UK) transfection reagent. Transfection media was added to cells in fresh SiHa or F-media without antibiotics. Twenty-four hours later, half of the transfection media was replaced with fresh growth media. Functional assays and RNA extractions were performed 48h later, while protein for western blotting was extracted 72h after transfection. siRNA information is listed in Supplementary Table 1.

### RNA isolation, cDNA synthesis, and quantitative PCR (qPCR)

Total RNA was extracted using the PureLink™ RNA Mini Kit (ThermoFisher) as per the manufacturer’s protocol, including an on-column DNase treatment. Purity and concentration of isolated total RNA were assessed using the NanoDrop® ND-1000 spectrophotometer (ThermoFisher). Complementary DNA (cDNA) was synthesized using the High-Capacity RNA-to-cDNA Kit (Applied Biosystems) and gene expression assessed using iTaq Universal SYBR Green Supermix (Bio-Rad, Hercules, CA) on a StepOnePlus Real-Time PCR System (Applied Biosystems). The thermal cycling protocol included an enzyme activation step at 94⍰°C for 2⍰min, followed by 40 cycles of a 15 ⍰s denaturing step at 94 ⍰°C, and a 1⍰min annealing/extension step at 55-60°C. Fluorescent intensity was measured at 65°C at the end of each cycle. Sequences of qPCR primers are listed in Supplementary Table 2.

### Analysis of single-cell RNA sequencing data

Two publicly available single-cell transcriptomic datasets from cervical squamous cell carcinoma were analyzed to characterize *TAOK3* expression and co-expression. The Fan et al. dataset (14) comprised 163,880 cells from 14 treatment-naive cervical squamous cell carcinoma tumors and 3 healthy donor cervix samples, annotated across 14 cell types. This dataset was available as a pre-processed Seurat object containing log1p-normalized expression values but no raw counts. Healthy donor samples were excluded, and analyses were restricted to tumor-derived samples. The Terekhanova et al. dataset (15) was obtained from the NCI Human Tumor Atlas Network pan-cancer single-nucleus atlas. Cervical squamous cell carcinoma tumor samples were extracted, comprising 14 patient samples with scRNA-seq data processed using CellRanger against GRCh38. Because raw counts were available, this dataset enabled additional count-based co-expression analysis. To balance representation, up to 600 tumor cells per sample were downsampled and merged into a single Seurat object. Analyses were performed in R v4.2.2 using Seurat v5 (16).

In the Fan et al. dataset, cells were restricted to Squamous and Glandular epithelial populations based on the cell_type_l1 annotation. In the Terekhanova et al. dataset, all retained cells were tumor cells by design, and no further cell type filtering was applied. No gene filtering was applied to the Fan et al. dataset before co-expression analysis. For the Terekhanova et al. dataset, Ensembl (RRID:SCR_002344) GRCh38 annotation via biomaRt (17) was used to retain protein-coding genes. Additional pattern-based filtering removed mitochondrial, ribosomal, unannotated, microRNA, small nucleolar RNA, and long non-coding RNA features, including *MALAT1, NEAT1*, and *XIST*. *TAOK3* was retained regardless of filtering status. *TAOK3* expression was assessed using log1p-normalized expression values. Highly variable genes were identified within each analysis subset using Seurat’s variance-stabilizing transformation method, selecting the top 5,000 features. *TAOK3* was added if absent, and the resulting feature set was scaled using ScaleData. Co-expression analyses were then performed in parallel using both log1p-normalized and scaled expression layers.

*TAOK3* co-expression was assessed using Pearson, Spearman, and Kendall correlations. Pearson measured linear association, Spearman measured rank-based association, and Kendall provided a concordance-based measure robust to non-normality and outliers. For Pearson and Spearman analyses, only *TAOK3*-expressing cells were included. Kendall correlation was additionally limited to a random maximum of 2,000 cells per analysis subset to reduce computational burden. Genes were ranked by their correlation coefficient with *TAOK3*. For the Terekhanova et al. dataset, CS-CORE (18) method was additionally applied using raw counts, which were unavailable for the Fan et al. dataset. CS-CORE was run on the top 2,000 protein coding genes by mean expression. When more than 5,000 cells were available, cells were randomly downsampled before analysis. *TAOK3* co-expression estimates were extracted, and p-values were adjusted using the Benjamini-Hochberg procedure; adjusted p-value < 0.05 was considered significant. Consensus *TAOK3* co-expressed genes in the Fan et al. dataset were defined as genes appearing among the top 20 ranked genes across Pearson, Spearman, and Kendall methods for a given analysis stratum and expression layer. In the Terekhanova et al. dataset, consensus support was summarized across Pearson, Spearman, Kendall, and CS-CORE, and visualized by tile-based heatmaps indicating method-level support.

### RNA sequencing (RNAseq)

Total RNA was extracted as described above and shipped on dry ice overnight to Novogene Co., LTD (Sacramento, CA) for library preparation and 150 bp, paired-end RNA sequencing on the Illumina NovaSeq 6000. Cells were passaged 3 times, and 2 replicate dishes were used for 4 total replicate samples (2 technical replicates per passage, 2 biological replicates; Supplementary Fig. S2A) except for MCW-3 (3 technical replicates, 1 biological). Raw sequencing reads were evaluated for quality control using FastQC v0.11.9 (https://www.bioinformatics.babraham.ac.uk/projects/fastqc/; RRID:SCR_014583). Metrics assessed included adapter content, read length distribution, and per-base sequence quality scores. A hybrid human-virus reference genome was created by concatenating the human primary assembly (GRCh38) with HPV 16 and 18 (Refseq ID: GCF_000863945.3 and GCF_003180735.1). Reads were aligned to the custom, hybrid human-virus reference genome using the STAR v2.7.10b aligner (RRID:SCR_004463; (19)) and transcript/gene-level expression abundance was quantified using RSEM v1.3.3 (RRID:SCR_013027; (20)). Genes with raw counts >10 in at least 3 samples were considered expressed and retained for further analysis. To assess sample similarity, Poisson Distance was computed using the CRAN package PoiClaClu version 1.0.2.1 (https://CRAN.R-project.org/package=PoiClaClu; RRID:SCR_003005). Principal Component Analysis (PCA) plots were generated using rlog-transformed values to visualize group relationships (Supplementary Fig. S2B). Differential expression (DE) analysis was performed using the Bioconductor (RRID:SCR_006442) package DESeq2 version 1.36.0 (RRID:SCR_000154; (21)) to compute log2FoldChange (L2FC) and False Discovery Rate (FDR) adjusted p-values. Statistical significance was determined at an FDR threshold of 0.05. Enrichment analysis of differentially expressed genes (DEGs; adjusted *p*<0.05; |L2FC| ≥ 0.58) was performed using Metascape (RRID:SCR_016620; (22)) or g:Profiler (RRID:SCR_006809; (23)).

### Proteomics and phosphoproteomics analyses

Cells (1.0 × 10^6^) were seeded in 10 cm dishes, allowed to adhere overnight, and then transfected with either siCONT or siTAOK3 siRNAs as described above. Cells were harvested 48 h later by washing with PBS, trypsinization, and centrifugation at 300 g for 5 min. Cell pellets were resuspended in PBS, washed again, and stored at – 80 °C until further use. Each sample was resuspended in 100 µL of 100 mM ammonium bicarbonate, 40% Invitrosol, and 20% acetonitrile, transferred to a 1.5 mL tube, and sonicated on ice for 25 cycles of 10 sec on and 30 sec off. Cysteines were reduced in 5 mM TCEP for 30 min at 37 °C, then alkylated with 10 mM iodoacetamide for 30 min at 37 °C. Samples were next digested overnight with 10 µg trypsin at 37 °C. After cleanup with PreOmics® Phoenix columns, peptides were reconstituted in 100 mM triethylammonium bicarbonate (TEAB) buffer, and concentration was measured using the Pierce Quantitative Fluorometric Peptide Assay. Each sample (100 µg) was labeled with TMT6plex reagents (ThermoFisher) following the manufacturer’s protocol. After labeling, samples were pooled and divided into an aliquot of 8.3% of the total, expected to contain 50 µg of labeled material. This aliquot was then dried, and the remainder was cleaned up using the SP2 method and dried. Dried samples were dissolved in 40 µL of 0.1% acetonitrile/0.1% formic acid for three 10 µL injections. For total proteomics analyses, one 50 µg aliquot of labeled sample was dissolved in 300 µLof 0.1% trifluoroacetic acid and fractionated using the Pierce High pH Reversed-Phase Peptide Fractionation Kit, using one water and one 5% acetonitrile wash, then eluting in 10, 12.5, 15, 17.5, 20, 22.5, 25, and 50% acetonitrile/TEA steps. Fractions were concatenated by combining 1+5, 2+6, 3+7, and 4+8 before drying and were redissolved in 50 µL 2% acetonitrile / 0.1% formic acid, diluted 5-fold further with Fisher Scientific Pierce Peptide Retention Time Calibration Mixture (4 nM).

For phosphoproteomics analyses, one 50 µg aliquot of the labeled sample was dissolved in 150 µL of binding/equilibration buffer and used for phosphopeptide enrichment with the Pierce High-Select TiO2 kit, as per the manufacturer’s protocol. The eluted material was dried, and the unbound and wash fractions were combined and dried. Unbound material was redissolved in 200µL of binding/equilibration buffer (Pierce High-Select Fe-NTA kit) and processed according to the manufacturer’s protocol, and the eluted material was dried. The peptides from the total proteomics and phosphoproteomics analyses were analyzed on a Thermo Scientific Orbitrap Fusion Lumos mass spectrometer. Samples were loaded onto a Thermo Acclaim PepMap C_18_ analytical column (75 µm x 50 cm) and resolved by applying a gradient of 2.5–9%, 9–34%, 34–60%, and 60–99% acetonitrile/0.1% formic acid at a flow rate of 300 nL/min. The mass spectrometer was operated in a data-dependent acquisition (DDA) HCD MS2 instrument method with a 3 s cycle time. The MS1 data were acquired over a scan range of 375–1500 *m/z* at 120,000 resolution, with a 200 *m/z* centroid. The precursor ions were isolated with an isolation window of 0.7 *m/z*, fragmented using HCD, and MS2 spectra were detected in the Orbitrap at 30,000 resolution at 200 *m/z*.

Raw mass spectrometry (MS) data were analyzed using MaxQuant (RRID:SCR_014485; (24)), employing the default search engine. Searches were performed against the UniProt human proteome database (RRID:SCR_002380; (25)) with key search parameters including: Enzyme specificity: Trypsin/P with up to 2 missed cleavages, Fixed modifications: Carbamidomethylation (C), TMT (K and peptide N-terminus), Variable modifications: Oxidation (M), Phosphorylation (S, T, Y), Mass tolerances: 20 ppm for first search, 4.5 ppm for main search, and FDR: <1% at peptide-spectrum match (PSM), protein, and site levels. Reporter ion intensities were corrected using the manufacturer-provided isotopic impurity tables. Phospho-site localization was assessed using MaxQuant’s built-in phospho (STY) site algorithm, and only sites with a localization probability >0.75 were considered for downstream analysis. MaxQuant output files (e.g., Phospho (STY)Sites.txt, proteinGroups.txt) were processed using Perseus (26). Reporter ion intensities were log2-transformed and normalized by subtracting the median across samples. Phospho-sites and proteins with valid values in at least 70% of samples in one group were retained. Missing values were imputed from a normal distribution to simulate low-abundance peptides. Differential analysis between experimental conditions was performed using two-sample Student’s t-tests with permutation-based FDR correction or Benjamini-Hochberg adjusted p-values, as appropriate. Significantly regulated proteins or phospho-sites were defined as those with |L2FC| ≥ 0.58 and adjusted p-value < 0.05. Data visualization was performed in Perseus (26) and pathway enrichment analysis and protein network generation were performed as listed above for RNAseq results. Furthermore, phosphoproteomics data were uploaded to the KSEA app to identify upstream master kinases (27).

### Histology

Sample preparation, processing, and staining were performed by the histology cores at Wisconsin Diagnostic Laboratories (human tissue; Leica Bond Prime automated stainer) or Children’s Research Institute (mouse tissue; Leica Bond Rx automated staining platform). Tissue/organoid samples were fixed in neutral buffered formalin (10%) and processed into paraffin blocks using a Tissue Tek VIP5 automated processor (Sakura, Japan). Paraffin blocks were sectioned at 4 µm thickness (Micron HM355S, Germany) and air-dried before staining. Routine hematoxylin and eosin (H&E) staining was performed on a Prisma Automated Stainer (Sakura, Japan). Briefly, sections underwent standard deparaffinization and rehydration, followed by hematoxylin staining (Richard Allen, 7211), differentiation in acid alcohol, and bluing with a 0.1% ammonia solution. Slides were then counterstained with eosin-Y (Richard Allen, 71211), differentiated in 100% ethanol and cleared in xylene. Coverslips were applied with synthetic mounting media and sections were imaged on a digital slide scanner (Hamamatsu).

### Immunohistochemistry/Immunofluorescence/Immunocytochemistry

Immunohistochemical (IHC) and immunofluorescent (IF) staining was performed on a Leica Bond Rx automated staining platform. Deparaffinization and antigen retrieval (Leica, DS9640) were performed before staining. Sections were blocked with Biocare Blocking solution (BS699) before antibody incubation. Primary antibodies were incubated for 1 hour at room temperature. Detection was achieved using either an HRP-tagged (*TAOK3*) or fluorophore-conjugated secondary antibody (CD206 or F4/80; Donkey anti-Rabbit Cy3, Jackson Immuno; 711-166-152) applied for 45 minutes at room temperature. Nuclei were counterstained with DAPI (Sigma, D8417; 1:5000), and sections were mounted with Prolong Gold antifade reagent (Invitrogen, P36930). Negative controls were generated by omission of the primary antibody. An IF of human tissue was imaged on a VS120 slide-scanning system (Olympus), and quantification and analysis were performed using the open-source software, QuPath (28), with the experimenter blinded to treatment group.

Immunocytochemistry was performed by washing 2D or 3D cells with PBS 3 times, incubating with permeabilization buffer (1% BSA and 0.1% Tween-20 in PBS), and then blocking at room temperature for 1 h in blocking buffer (PBS + 10% BSA, 0% normal donkey serum, and 0.1% Tween-20). Primary antibodies in the blocking buffer were incubated overnight at 4°C. The next day, cells were again washed 3 times with PBS, then incubated with fluorescent secondary antibodies for 1 h at room temperature. After a final PBS wash, cells were covered with an antifade media for imaging. All antibodies are listed in Supplementary Table 3. Confocal images were acquired on a Andor BC43 benchtop spinning disk confocal located in the MCW Oxford Center for Advanced Microscopy – Electron Microscopy Core (OxCAM-EM).

### Incucyte cell cycle analysis

SiHa cells were transduced with the Incucyte Cell® Cycle Lentivirus reagent (Sartorius; #4779) according to manufacturer’s instructions. Briefly, cells (2000/well) were seeded in a 96-well plate. Once adhered, the cell cycle lentivirus reagent was added to the media supplemented with polybrene (8 µg/ml). Twenty-four hours later, the media was replaced with fresh growth media, followed by puromycin (0.1 µg/ml) selection. The final puromycin-resistant cell population remaining was grown up and labeled Incucyte SiHa (Inc-SiHa). Inc-SiHa cells were seeded and grown for 24 h, after which thymidine was added to a final concentration of 2.5 mM, and cells were incubated for 24 h (first block). Cells were released for 12 h in complete medium, then blocked again in 2.5 mM thymidine for 24 h. The final release of the cells was performed in complete medium for varying lengths of time to examine alterations in cell cycle dynamics in *TAOK3* inhibitor-treated cells (SBI-581; 100 nM) versus control-treated cells (DMSO). Cells were subjected to automated imaging and quantitative analysis using an Incucyte® S3 live imager.

### Cell growth assays

Colony formation assays were performed by seeding transfected cells (2000/well) into 6-well plates and incubating in a humidified, 5% CO_2_ atmosphere. Fresh growth media were replaced every 3-4 days and colonies were fixed and stained with crystal violet (Sigma) after 2 weeks *in vitro*. Colony assay imaging and analysis were performed on the iBright Imaging System (ThermoFisher). Cell proliferation was measured using the CellTiter 96® AQueous One Solution Cell Proliferation Assay (Promega, Madison, WI) according to the manufacturer’s directions. Cells (transfected or untransfected for inhibitor studies) were seeded onto 96-well plates (100⍰µl of 0.02⍰×⍰10^6^ cells/ml per well) and allowed to adhere overnight. For assays using the *TAOK3* inhibitor (SBI-581; (29,30)), cells were incubated with the inhibitor overnight before and throughout chemotherapy. MCW-3 spheroids were generated by seeding 5000 cells/well in an ultra-low attachment 96-well plate. Spheroids were grown *in vitro* for 72hr before addition of DMSO or SBI-581 (10 nM) to the culture media. Following this overnight pretreatment, cells were treated with DMSO or one of two taxol doses (1 nM or 100 pM) with or without SBI-581 for an additional 72 hr. Cell viability was then assessed using the CellTiter-Glo® 3D Cell Viability Assay (Promega), according to the manufacturer’s protocol. Briefly, an equal amount of CellTiter-Glo® 3D Reagent was added to each well containing a single MCW-3 spheroid and mixed by vigorous shaking for 5 minutes. The spheroids were then incubated at room temperature for another 25-30 min before recording luminescence on a microplate reader (Tecan Spark, Switzerland).

### Western blotting

Monolayer cells (~3 × 10^6^) were washed with 1x PBS and incubated in RIPA cell extraction buffer (ThermoFisher) supplemented with 1X Halt Protease and Phosphatase inhibitor cocktail (ThermoFisher) at −20°C overnight. The next day, protein lysates were thawed on ice and homogenized by running through a 21G needle. Undigested cellular debris was pelleted by centrifugation at 10,000 rpm for 5 min at 4°C and protein was quantified using a standard Bradford Assay (Sigma). Cell lysate (30 μg) was run on a Criterion Tris-HCl polyacrylamide gel (Bio-Rad), transferred to PVDF membranes, blocked with 1x TBS + 10% w/v nonfat dry milk at room temperature for 1h, and incubated with primary antibody overnight at 4°C. The following day, membranes were washed, incubated for 1h at room temperature with HRP-conjugated goat anti-rabbit or mouse anti-rabbit IgG (Cell Signaling Technology; Danvers, MA), and developed using ECL (Life Technologies). Chemiluminescence was visualized using an iBright Imaging System (ThermoFisher) and the visualized protein bands were quantified using the iBright imaging software. Primary antibodies used are listed in Supplementary Table 3.

### Generation and validation of stable CRISPRi SiHa cell lines

The pHR-UCOE-EF1a-Zim3-dCas9-P2A-BFP plasmid was a gift from Marco Jost and Jonathan Weissman (Addgene plasmid # 188777; http://n2t.net/addgene:188777; RRID:Addgene_188777; (31)). The plasmid was packaged into lentivirus by the Viral Vector Core Lab at the Versiti Blood Research Institute following standard protocols. Parental SiHa cells were stably transduced with the pHR-UCOE-EF1a-Zim3-dCas9-P2A-BFP lentiviral construct, in the presence of 8 µg/ml polybrene, following the Weissman lab protocol ((31); 5 MOI). Following 48-72 h transduction, polyclonal populations of BFP-positive cells were selected using 2 rounds of FAC sorting on a BD FACSAria III, with FACSDiva software for analysis. Cells were sorted into FBS and then transferred to appropriately sized plates containing media consisting of DMEM, Penicillin-Streptomycin (1%), Amphotericin B (25 µg/ml) and additional FBS (20% v/v). Cells were maintained in media containing antifungal, antibiotics, and extra FBS for ~2 weeks before being transitioned back to regular growth media.

### Design, cloning, and transduction of dual guide RNAs

Non-targeting control (sgNTC) and *TAOK3*-targeting (sgTAOK3) dual sgRNA oligos from Replogle et al., 2022 (31) were generated by IDT (Integrated DNA Technologies, Inc.) and cloned into pJR104 (Addgene #187243) as described ((31); https://weissman.wi.mit.edu/resources/). Briefly, pJR104 vector and annealed dual sgRNAs were digested with Blpl and BstXI, followed by ligation. Next, the sgRNA CR3/hU6 promoter from pJR98 (Addgene #187239) was BsmBI-digested and inserted into the dual sgRNA-containing pJR104 to ensure both sgRNAs have a U6 promoter and constant region (CR). After Sanger sequencing validation (Functional Biosciences, Madison, WI), resultant plasmids were packaged into lentivirus by the Viral Vector Core Lab at the Versiti Blood Research Institute (VBRI, Milwaukee, WI) following standard protocols. Stable CRISPRi SiHa cells were seeded (2.0⍰×⍰10^5^) and then transduced 24 h later with sgNTC or sgTAOK3 (5 MOI) with 8 µg/ml polybrene. Seventy-two hours later, cells were sorted to select for those expressing both BFP and GFP. Cells were sorted into FBS and then transferred to appropriately sized plates containing media consisting of DMEM, Penicillin-Streptomycin (1%), Amphotericin B (25 µg/ml) and additional FBS (20%). Cells were kept in media containing antifungals and antibiotics, with extra FBS, for 2 weeks, then transitioned back to DMEM + 10% v/v FBS. Western blot and qPCR were performed to confirm CRISPRi-mediated *TAOK3* knockdown. Sequences of dual sgRNAs are listed in Supplementary Table 1.

### *in vivo* experiments

For *in vivo* experimental purposes, 4–6-week-old female athymic nude mice (Athymic Nude-Foxn1nu) were purchased from Envigo and housed in pathogen-free conditions. All experiments on mice were performed in accordance with the Medical College of Wisconsin Institutional Animal Care and Use Committee (IACUC). Animal health was monitored daily. For orthotopic tumor cell inoculation, control (sgNTC) or *TAOK3* knockdown (sgTAOK3) CRISPRi SiHa cells were trypsinized, washed, and resuspended in Ca^2+^ and Mg^2+^-free 1x Hanks’ balanced salt solution (HBSS, GIBCO, Carlsbad, CA, USA), and 3×10^6^ cells were injected into the cervix of mice (n=10/group). Tumor growth was monitored weekly by bioluminescence imaging (IVIS). At the end of the experiment, the mice were sacrificed, and tumors were dissected, weighed, and processed for downstream applications.

### Species-specific RNA sequencing (SSRS)

RNA sequencing was performed on the Illumina NovaSeq X Plus platform using paired-end 150 bp (PE150) chemistry. Libraries were sequenced to sufficient depth to enable comprehensive transcriptome profiling and differential expression analysis. On average, 1.71 × 10^8^ ± 4.28 × 10^7^ reads per sample were generated. Raw sequencing reads were quality-controlled using FastQC v0.11.9 (http://www.bioinformatics.babraham.ac.uk/projects/fastqc) and processed with Trimmomatic v0.38 (RRID:SCR_011848; (32)) to remove adapters and low-quality bases. To enable simultaneous detection of human, mouse, and HPV16 transcripts, we constructed a comprehensive multi-species reference genome. The human reference genome (GRCh38.p14, GENCODE release 44; RRID:SCR_014966) and mouse reference genome (GRCm39, GENCODE release M33) were obtained from the GENCODE database. The HPV16 reference genome (NC_001526.4) was retrieved from NCBI. To prevent cross-species mapping artifacts, chromosome and contig identifiers were prefixed with species-specific tags (“human_”, “mouse_”, and “HPV16_” respectively) before concatenation into a single reference file. Corresponding GTF annotation files were downloaded from GENCODE, and the HPV16 genome was downloaded from NCBI, encompassing all annotated transcripts, including protein-coding genes, long non-coding RNAs, and pseudogenes. Quality-filtered reads were then aligned to the multi-species reference genome using the STAR v2.7.10b aligner (19), followed by gene expression quantification using RSEM v1.3.3 (20). Species-specific expression profiles were separated by gene identifier prefix (human: ENSG; mouse: ENSMUSG; HPV16: HPV16/hpv16), with transcripts per million (TPM) values calculated for each species and normalized to total TPM across all three genomes (Supplementary Table 4). Differential expression analysis was performed on xenograft samples (*n* =4 control, *n* =4 TAOK3 KD) using DESeq2 v1.42.1 (21), with significantly differentially expressed genes defined as having adjusted p<0.05 and |L2FC| >0.58.

### THP-1 co-culture and Incucyte imaging/analysis

THP-1 cells (ATCC #TIB-202) were grown at 5% CO_2_, 37°C in RPMI 1640 Medium (ATCC modification; ThermoFisher #A1049101). THP-1 cells were seeded into a 24-well plate at 10^5^ cells/well and treated with 100ng/mL phorbol 12-myristate 13-acetate (PMA; Sigma-Aldrich P1585) for ~24h. PMA-containing media was replaced with fresh media for another 24h before adding 10^5^ SiHa sgNTC or SiHa sgTAOK3 cells to the THP-1 cultures. Plates were placed into an Incucyte® S3 live imager and scanned every 30 min for 72h to capture brightfield and GFP images. Image analysis was performed using Incucyte software. Macrophages were defined based on area (300 – 950um^2^), eccentricity (maximum 0.95) and green mean intensity (maximum of 0.1500), and their eccentricity and size were plotted over time after normalizing to time 0.

### Lucifer yellow uptake and Lysotracker staining

Transfected cells were seeded in a 96-well plate (5000 cells/well) and allowed to adhere overnight. Medium was replaced with media containing lucifer yellow (LY; ThermoFisher; 0.5 mg/ml) and either DMSO, taxol (1 nM), or the methuosis inducer, MOMIPP (10 µM). Plates were then immediately placed in the Incucyte® S3 live imager and set to capture brightfield and LY (green channel) images every hour for 24h. The next day, Lysotracker Deep Red (ThermoFisher; 75 nM) was added to each well, and wells were scanned on all 3 channels (brightfield, green, and red) at 15 min intervals for 2-4 h. Images were analyzed using the Incucyte basic analyzer software to calculate amount of colocalization of lucifer yellow-filled cytoplasmic vacuoles with lysosomes (non-colocalized green = total green – overlapping green (yellow)).

### Extracellular ATP assay

Extracellular ATP was measured using the RealTime-Glo™ Extracellular ATP Assay (Promega) according to the manufacturer’s instructions. Briefly, transfected cells were seeded (5000 cells/well) in a 96-well plate and allowed to attach overnight. The following day, the RealTime-Glo™ Extracellular ATP Assay reagent was added to each well and luminescence was measured over 24 h using a Tecan Spark multimodal plate reader.

### Data Availability

RNA sequencing data analyzed in this study are available in the NCBI Sequence Read Archive (SRA) under submission SUB16221097.

### Statistical analysis

Functional assays were performed in triplicate. All statistical analyses and graphing of data were performed using GraphPad Prism 11.0.1 (RRID:SCR_002798; La Jolla, CA).

## Results

### *TAOK3* expression is upregulated in cervical cancer and localized to a distinct subpopulation of tumor epithelial cells

We first investigated *TAOK3* expression level and localization in our banked cervical cancer tissue and cell lines (**Fig. 1)**. *TAOK3* mRNA was elevated in cervical tumor tissue (*n* =131) compared to adjacent normal (*n* =9; *p* =0.086; **Fig. 1A)**. RNA sequencing (RNAseq) and western blotting of commercially available (SiHa) and patient-derived cervical cancer cell lines (MCW-1, MCW-2, and MCW-3) confirmed *TAOK3* mRNA (**Fig. 1B)** and protein (**Fig. 1C)** expression, respectively. TAOK3 protein was primarily localized to the proliferating outer edges of 3D SiHa and MCW-3 spheroids and exhibited colocalization with the leader cell marker KRT14 (**Fig. 1D)**. Western blotting confirmed *TAOK3* expression in cytoplasmic and membrane compartments of these 2D cultured cells (**Supplementary Fig. S1A)**. Further, we observed overlapping *TAOK3* expression with HPV mRNA expression in patient-derived tumoroids (**Fig. 1E and Supplementary Fig. S1B)** and higher TAOK3 expression in cervical cancer tumor and metastases compared to normal cervix by IHC (**Fig. 1F and Supplementary Fig. S1D)**. Although TAOK3+ staining was observed in keratin-expressing tumor epithelial cells (**Supplementary Fig. S1C)**, it was not always ubiquitous. Thus, we leveraged data from two cervical cancer single-cell RNAseq (scRNA-seq) studies (14,15) to elucidate the transcriptomic signature of TAOK3+ cancer epithelial (T3epi) cells. We investigated cell clustering (**Fig. 1G and I)** and single-cell *TAOK3* expression (**Fig. 1H and J)** for each dataset and then bioinformatically identified co-expressed genes in T3epi cells (**Supplementary Tables 5 and 6)**. Gene Enrichment analysis (g:Profiler;(23); **Supplementary Fig. S1E and Table 7)** of genes expressed in T3epi cells (n = 84 co-expressed genes across both datasets with 3 overlapping) revealed enrichment for cadherin binding (*p* =0.0004), S100 protein binding (*p* = 0.011), epithelial cell differentiation (*p* = 0.038), plasma membrane (*p* =1.56E-05), vesicles (*p* = 0.0002), and cell leading edge (*p* =0.046). Of note, three genes were found in both datasets (*EPS8L1, RNF213*, and *SPNS2*). Collectively, these data provide evidence for a subpopulation of keratin-expressing T3epi cell population positioned at the invasive front and enriched at membrane/actin interfaces, consistent with a pro-migratory, leader cell–like state.

**Figure 1.**
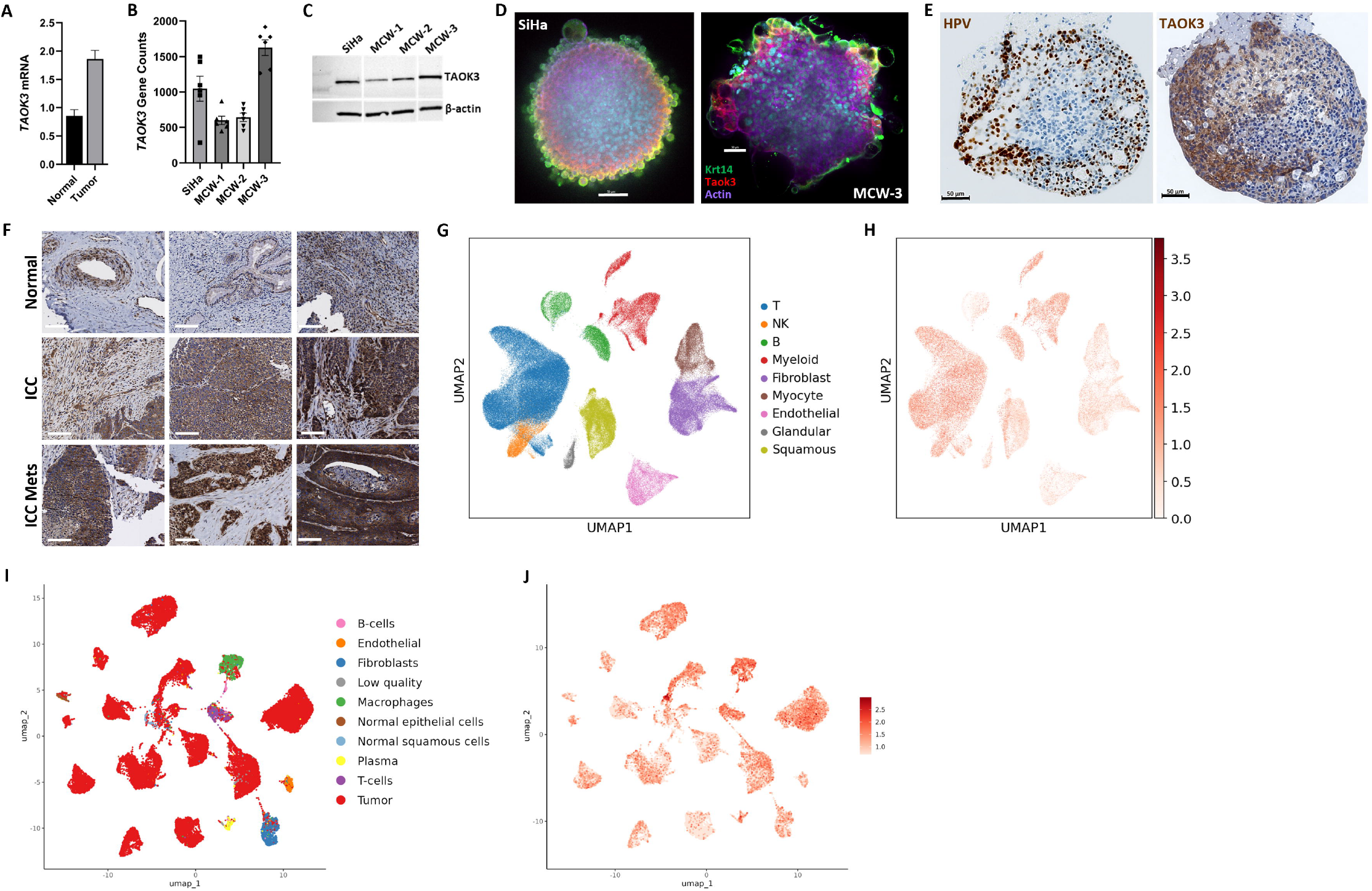
*TAOK3* is expressed in a distinct subpopulation of cervical cancer epithelial cells. **A**, *TAOK3* mRNA expression is higher in cervical cancer tumor tissue compared to adjacent normal (*p* =0.86; *n* = 9 normal and *n* = 131 tumor). **B**, *TAOK3* mRNA and **C**, protein expression in 4 unique cervical cancer cell lines. **D**, *TAOK3* (red) protein expression is colocalized with keratin (KRT14 in green) at the outer edges of 3D Si Ha and MCW-3 spheroids. Scale bar (white)= 50 µm. **E**, *TAOK3* protein expression overlaps HPV+ cells in a patient-derived cervical cancer tumoroid. Scale bar (black)= 50 µm. **F**, *TAOK3* IHC staining in normal cervix, invasive cervical cancer (ICC), and metastatic sites show increased staining in ICC and ICC Mets compared to normal. Scale bars (white)= 100 µm. G, UMAP plot showing the distribution of major cell types from the Fan et al., 2023 dataset. H, *TAOK3* single cell expression from Fan et al., 2023. **I**, UMAP plot showing the distribution of major cell types from the Terekhanova et al., 2023 dataset. **J**, *TAOK3* single cell expression from Terekhanova et al., 2023.

### *TAOK3* knockdown reprograms cervical cancer transcriptomes toward reduced cell cycle and motility with coordinated shifts in WNT, cytoskeletal, and immune pathways

To examine the functional role of T3epi cells in ICC, we employed siRNA-mediated *TAOK3* KD (siTAOK3) followed by RNA-seq analysis in 4 cell lines; **Supplementary Fig. S2A, 2B)**. Differential expression analysis (**Fig. 2A-D a**nd **Supplementary Table 8)** revealed cell line-specific variation in the number of differentially expressed genes (DEGs; p<0.05; |L2FC| cutoff of 1.0; FDR≤0.05; **Supplementary Fig. S2C)**, with 76 downregulated (**Supplementary Fig. S2D)** and 21 upregulated (**Supplementary Fig. S2E)** common across all four *TAOK3*-silenced cell lines. Metascape (22) analysis of the shared DEGs from all *TAOK3* KD cervical cancer cells showed enrichment in WNT signaling, mitotic cell cycle processes, negative regulation of protein metabolic processes, intrinsic apoptotic signaling in response to DNA damage, and regulation of vesicle-mediated transport (**Fig. 2E). Fig. 2F f**urther highlights expression of specific DEGs enriched in WNT signaling, cell cycle, motility and death programs, and tumor microenvironment (TME) pathways, some of which were further validated via qPCR (**Supplementary Fig. S2F)** and/or western blotting (**Supplementary Fig. S2G)**. When combining all significant DEGs from the four cell lines (*p*<0.05; |L2FC| ≥ 1.0; FDR≤0.05), downregulated genes were enriched for immune signaling and processes involving the actin cytoskeleton and movement (**Supplementary Fig. S2H)**. Upregulated DEGs were enriched for signaling by Rho GTPases, Miro GTPases, and RHOBTB3, MAPK cascade regulation, epithelial cell differentiation, Hippo Merlin signaling dysregulation, and endocytosis (**Supplementary Fig. S2I)**. Notably, transcripts associated with TME signaling and innate immunity were differentially regulated: pro-inflammatory cytokines/chemokines (e.g., *IL1A, IL6, CXCL1, CXCL8*) and myeloid/colony-stimulating factors (e.g., *CSF2/CSF3*) changed in a manner indicative of altered macrophage/immune crosstalk, while *DDX58* (RIG-I) was reduced across all cell lines. Together, these data indicate that *TAOK3* supports cell-cycle progression, WNT-driven motility programs, and immune/TME signaling and silencing of *TAOK3* redirects transcription toward a less proliferative, less migratory state with remodeled inflammatory cues.

**Figure 2.**
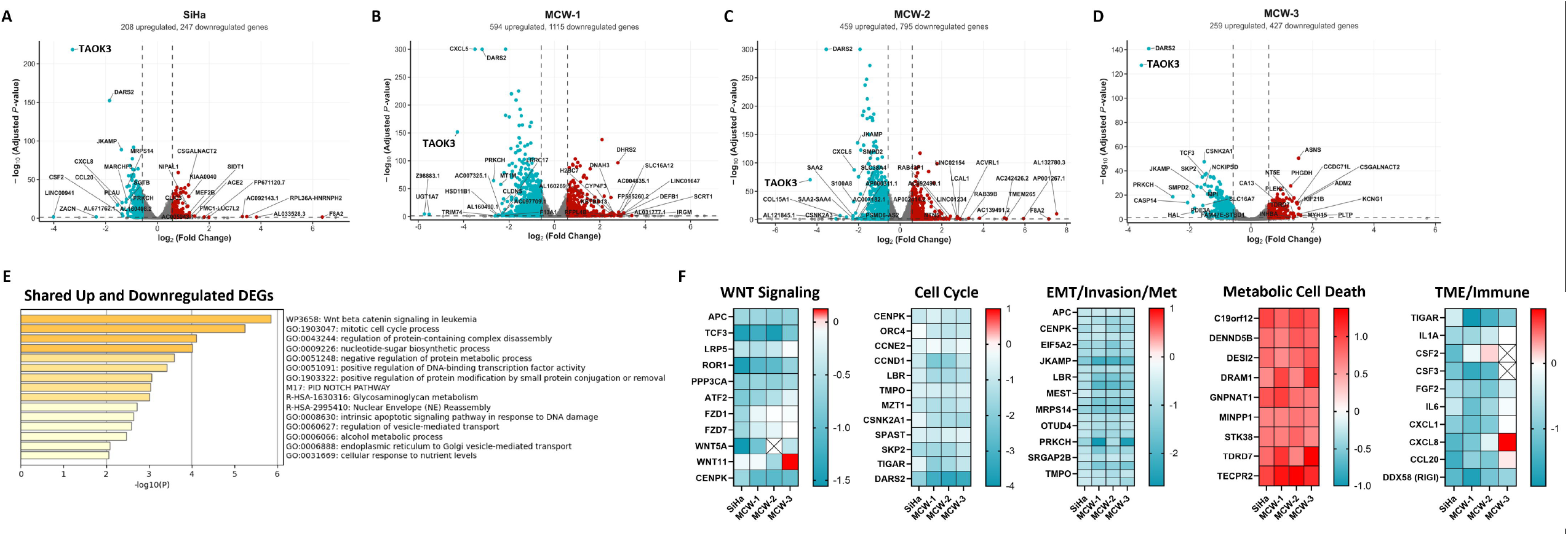
*TAOK3* knockdown reprograms cervical cancer transcriptomes toward reduced cell cycle and motility with coordinated shifts in WNT, cytoskeletal, and immune pathways. **A-D**, Volcano plots showing differentially expressed genes following siRNA-mediated *TAOK3* knockdown in four cervical cancer cell lines (*p*<0.01; L2FC= I0.58 I; FDR≤0.05). **E**, Metascape analysis of common DE genes across the four *TAOK3*-silenced cell lines (*n* =76 genes down and *n* = 21 genes up; *p*<0.05; L2FC2≥ |1.0|; FDR≤0.05) **F**, Heatmaps of DEGs by pathway in the four cervical cancer cell lines. Negative L2FC values from RNAseq data are in blue, while positive are in red. No color= no significant difference between siCONT and si*TAOK3*. Boxes with an X depict genes not expressed in associated cell line.

### *TAOK3* silencing remodels the tumor proteome and phospho-signaling networks governing endocytosis, cytoskeleton, and immune pathways

Next, we performed mass spectrometry on *TAOK3* KD cells to identify potential downstream effectors and roles of this kinase in cervical cancer (**Fig. 3)**. Total proteomics of *TAOK3*-silenced SiHa, MCW-2, and MCW-3 cells identified differentially expressed proteins (|L2FC| ≥ 0.58; 2.5% FDR; *p* ≤ 0.05; **Fig. 3A-C and Supplementary Fig. S3A)**. In SiHa, differentially expressed proteins were strongly enriched in antiviral innate immune response and positive regulation of leukocyte-mediated immunity (**Fig. 3D;** downregulated proteins: RIGI (DDX58), ATAD3A, IFIT2, and IFIT1; upregulated proteins: HLA-A and HLA-H). RIG-I (DDX58) mRNA downregulation was also observed in the transcriptomic data from *TAOK3*-silenced cell lines. Key pathways enriched in *TAOK3*-silenced MCW-2 were related to endocytosis, translation, and metabolism of RNA (**Fig. 3E)**, while differentially expressed proteins in *TAOK3*-silenced MCW-3 cells were enriched for regulation of exosome secretion, protein folding, and mitotic anaphase (**Fig. 3F)**. DARS2 (Aspartyl-TRNA Synthetase 2, Mitochondrial) mRNA was significantly downregulated in all four *TAOK3* KD cell lines and DARS2 protein was significantly downregulated in MCW-2 and MCW-3 KD cells. Proteins upregulated in more than one *TAOK3* KD cell line were the DNA damage repair nuclease FAN1 (FANCD2 and FANCI Associated Nuclease 1; SiHa and MCW-2) and the beta-tubulin folding regulator, RP2 (RP2 Activator of ARL3 GTPase; MCW-2 and MCW-3; (33)).

**Figure 3.**
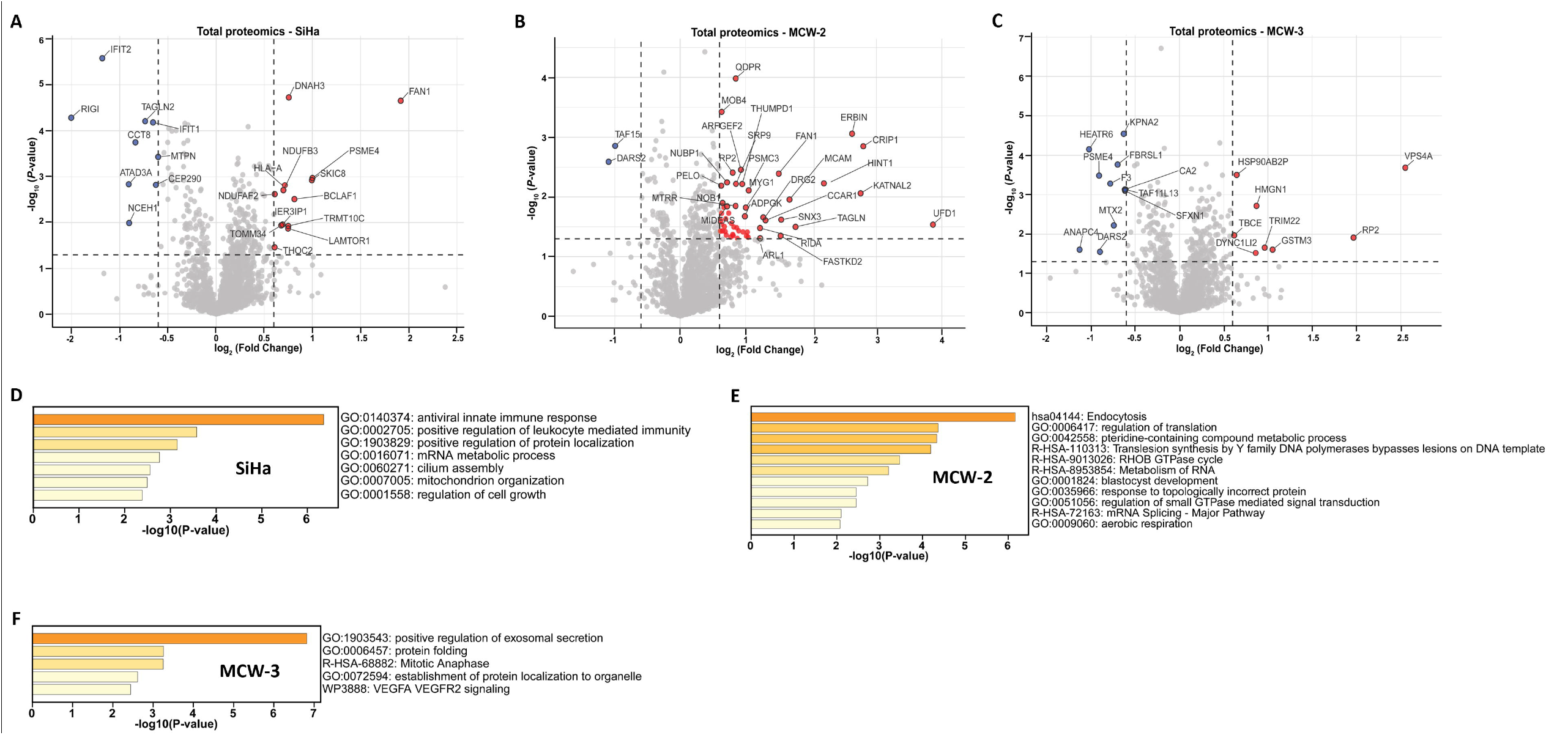
Proteomics profiling of *TAOK3*-silenced cervical cancer cell lines. **A-C**, Volcano plots depicting significantly altered protein expression in *TAOK3*-silenced SiHa, MCW-2, and MCW-3 cells, respectively. Blue dots= significantly downregulated proteins; red dots= significantly upregulated proteins. Metascape enrichment of significantly altered protein expression in TAOK3-silenced SiHa **(D)**, MCW-2 **(E)**, and MCW-3 **(F)**.

We next performed phosphoproteomics analysis of *TAOK3*-silenced SiHa, MCW-2, and MCW-3 cells (**Fig. 4 a**nd **Supplementary Fig. S3B)**. In SiHa cells, 89 phospho-sites in 74 different proteins were differentially phosphorylated after *TAOK3* knockdown (**Fig. 4A)**. MCW-2 and MCW-3 exhibited 39 phospho-sites from 33 different proteins and 100 phospho-sites from 88 different proteins, respectively (|L2FC| ≥ 0.58; 2.5% FDR; *p* ≤ 0.05; **Fig. 4B and 4C)**. Enrichment analysis of differentially phosphorylated sites in SiHa showed significant enrichment of mRNA metabolic process chromatin remodeling, Interferon signaling, JAK/STAT signaling, CDC42 GTPase cycle (and regulation of cytoskeleton organization pathways (**Supplementary Fig. S3C)**. MCW-2 phosphoproteomics data revealed enrichment chromatin remodeling, RAC1 GTPase cycle, ciliary landscape, endocytosis, and cell cycle (**Supplementary Fig. S3D)**. Finally, MCW-3 phosphoproteomics data were enriched for the following pathways: mRNA processing, chromatin remodeling, ciliary landscape, cell cycle process, regulation of nucleocytoplasmic transport −, and ERAD pathway (**Supplementary Fig. S3E)**. Finally, we performed Kinase–Substrate Enrichment Analysis (KSEA; (27)) of *TAOK3* KD phosphoproteomics data followed by gene ontology (GO) enrichment analysis to determine significant activity changes and delineate their biological implications (**Figs. 4D-F)**. Not many kinases showed increased activity, but all three TAOK3 KD cell lines showed decreased activity of kinases involved in cytoskeletal or cell-projection signaling, ERBB2-ERBB3 signaling, and transport regulation. Together, the proteome–phosphoproteome integration positions *TAOK3* as a coordinator of cytoskeleton–membrane dynamics and vesicular systems that interface with innate immunity and mitotic progression, providing a mechanistic basis for the previously reported defects in invasion, G2/M progression, and tumor–immune remodeling upon *TAOK3* inhibition.

**Figure 4.**
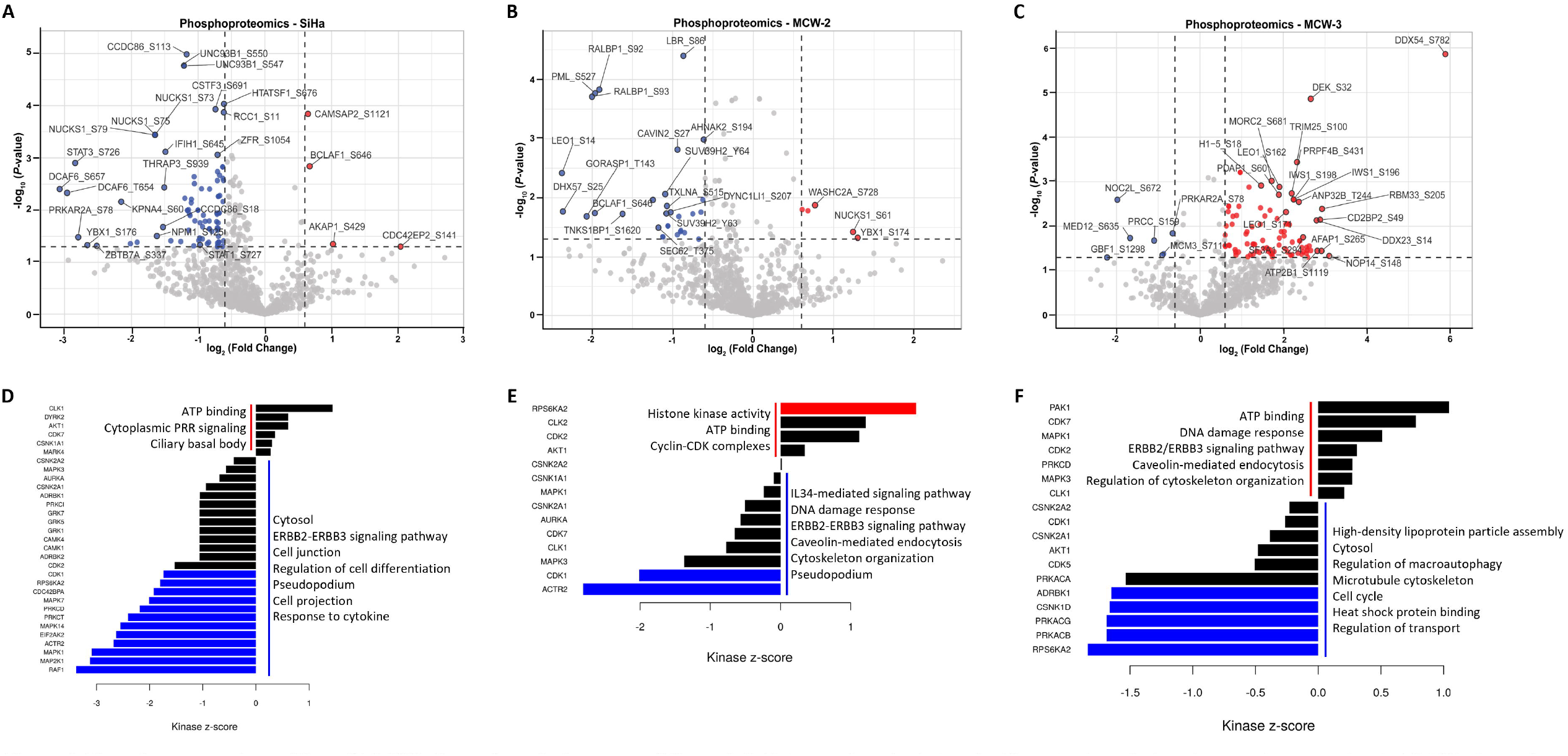
Phosphoproteomic profiling of *TAOK3*-silenced cervical cancer cell lines. **A-C**, Volcano plots depicting significantly altered phosphoprotein expression in *TAOK3*-silenced SiHa, MCW-2, and MCW-3 cells, respectively. Blue dots= significantly downregulated phosphoproteins; red dots= significantly upregulated phosphoproteins. Kinase-Substrate Enrichment Analysis (KSEA) of differentially expressed phosphoproteins in SiHa **(D)**, MCW-2 **(E)**, and MCW-3 **(F).** Blue= predicted to be significantly downregulated in activity; Red = predicted to be significantly upregulated in activity.

### *TAOK3* inhibition affects cell cycle, invasion, and chemoresponse *in vitro* and decreases tumor burden *in vivo*

Based on the transcriptomic and proteomic results, we next employed the Sartorius IncuCyte® system to examine cell cycle dynamics after *TAOK3* KD or inhibition. IncuCyte SiHa (Incu-SiHa) cells were generated using the Incucyte Cell Cycle Lentivirus and subjected to a double-thymidine block to synchronize cells in the G1/S phase and then released and monitored over the next 48 hours. **Figures 5A and 5B d**epict the percentage of total cells in each phase of the cell cycle following release from thymidine block in control DMSO or *TAOK3* inhibitor-treated (SBI-581 (29,30)) Incu-SiHa cells, respectively. SBI-581-treated Incu-SiHa cells exhibited a signifcantly prolonged G2M phase compared to DMSO-treated cells (**Fig. 5A and 5B;** green traces). Immediately following their release from the thymidine block (**Fig. 5C and Supplemental Fig. S4A;** 0 hr), SBI-581-treated cells had significantly fewer cells in the G1 phase than control cells. However, 18 hours after release (**Fig. 5C and Supplemental Fig. S4A;** 18h), we found significantly more *TAOK3* KD cells in the G2M phase than in control cells, resulting in a decreased growth rate (**Supplementary Fig. S4B)**. *TAOK3*-silenced cells also decreased the invasiveness of cervical cancer cells (**Fig. 5D and 5G; Supplementary Fig. S4C and S4D)**. Previous studies have shown that *TAOK3* inhibition can affect the response to anti-microtubule agents such as paclitaxel (taxol; (34)). Taxol is one of the most common chemotherapeutic agents used to treat cervical cancer, thus we next determined the effect of *TAOK3* KD/inhibition on taxol response in the two cervical cancer cell lines with the highest *TAOK3* expression, SiHa and MCW-3. We used sub-IC_50_ doses of taxol in combination with *TAOK3* KD and observed that, in the absense of *TAOK3* activity, cell proliferation in response to low-dose taxol is significantly decreased in *TAOK3* KD compared to control SiHa (**Fig. 5E)** and MCW-3 (**Fig. 5H)** cells. Pretreatment with SBI-581 (50 nM) in both SiHa (**Fig. 5F)** and MCW-3 (**Fig. 5I)** showed significantly decreased cell proliferation, specifically when paired with lower doses of taxol.

**Figure 5.**
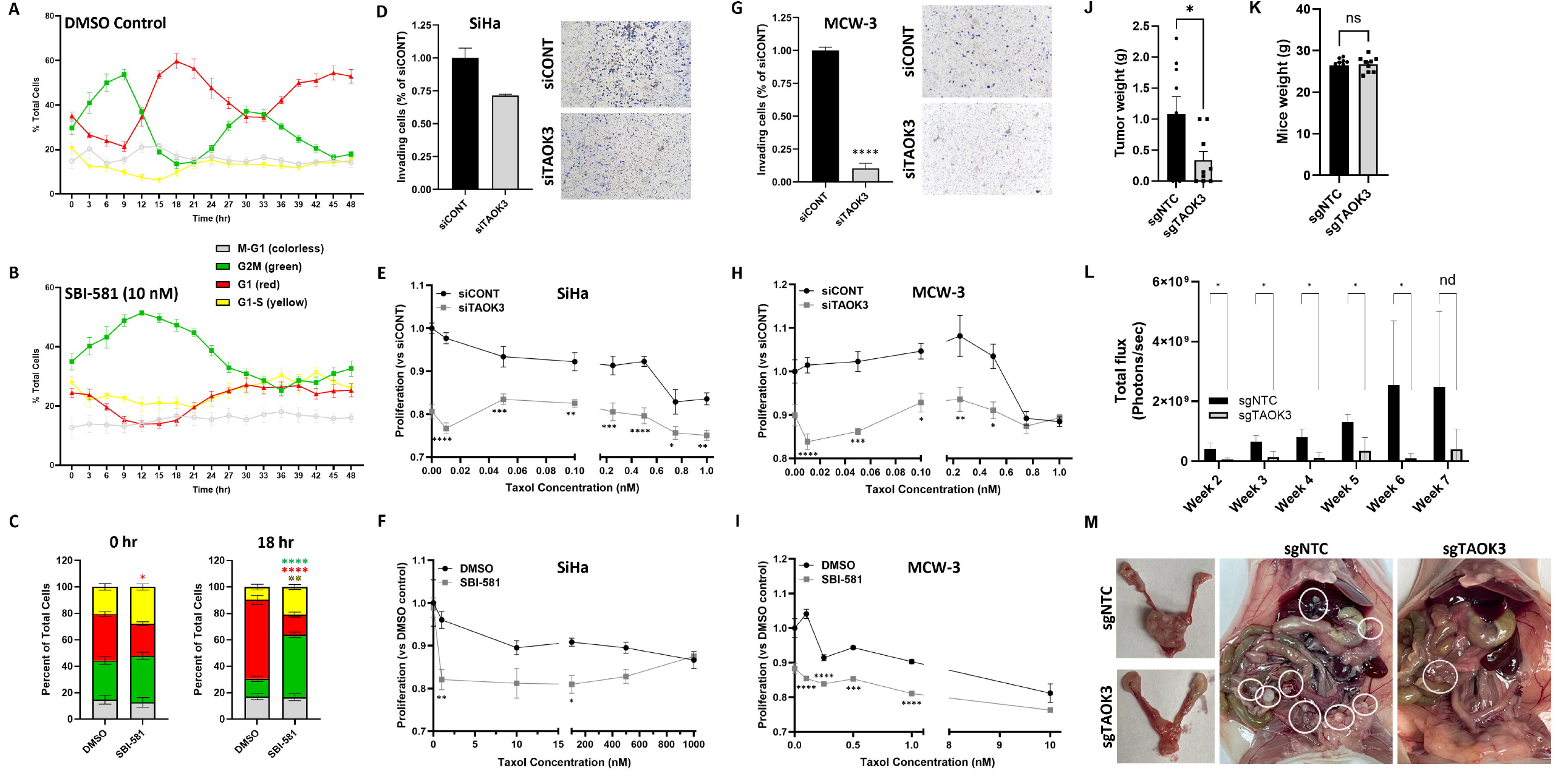
*TAOK3* inhibition affects cell cycle, invasion, and chemoresponse *in vitro* and decreases tumor burden *in vivo*. **A**, Percent of DMSO-treated lncuCyte SiHa (lncu-SiHa) cells in each phase of the cell cycle is shown across time. B, Same as in A but in SBl-581-treated lncu-SiHa cells. **C**, Bar graph depicting percent of total cells in different phases of the cell cycle in DMSO- or SBl-581-treated (50 nM) lncu-SiHa cells immediately following release from thymidine block (0 hr, left) and 18 hr after block released (right). **D**, siTAOK3 SiHa cells exhibited a trend (*p* =0.064) toward decreased invasion compared to siCONT SiHa. TAOK3 silenced **(E)** or inhibited **(F)** SiHa showed increased response to sublethal taxol doses. **G**, siTAOK3 MCW-3 cells exhibited significantly decreased invasion compared to siCONT MCW-3 cells. TAOK3 silenced **(H)** or inhibited (I) MCW-3 cells showed increased response to sublethal taxol doses. **J**, Weight of tumors from sgNTC mice was significantly higher than in tumors from mice injected with sgTAOK3 cells. **K**, Total body weight was not significantly different in sgNTC vs. sgTAOK3 injected mice. **L**, Bioluminescence imaging of mice across 7 weeks showed significantly less flux in sgTAOK3 vs. sgNTC mice, confirming decreased tumor burden in these animals. **M**, Gross images of extracted reproductive tracts (left panels) and abdomens (right two panels) from sgNTC and sgTAOK3 injected mice. White circles denote visible tumor nodules. *\*p<0.05; **p<0.0l; ***p<0.001; ****p<0.0001*.

Our previous work (5) showed that *TAOK3* KD in SiHa and HeLa cervical cancer cells significantly reduces colony formation and growth, so we next sought to evaluate *TAOK3*’s role in regulating cervical tumor progression in a mouse xenograft model. First, we generated CRISPRi-mediated *TAOK3* KD (sgTAOK3) and negative control (sgNTC) cells using dual sgRNAs as previously described (31). We verified the specificity and efficacy of CRISPRi-mediated *TAOK3* knockdown at the mRNA and protein level (**Supplementary Fig. S5A-B)** and observed that, like siRNA-mediated KD, sgTAOK3 cells exhibited significantly decreased spheroid diameter (**Supplementary Fig. S5C)** and colony number and area (**Supplementary Fig. S5D)** compared to sgNTC cells, *in vitro*. Next, firefly luciferase-labeled sgNTC and sgTAOK3 cells (3 × 10^6^) were injected intra-cervically into nude mice to determine the effect of TAOK3 KD *in vivo*. Seven weeks after injection, we observed no significant difference in overall body weight between sgTAOK3 and sgNTC-injected mice (**Fig. 5K)**. However, among sgTAOK3 mice that developed tumors (6 of 9 mice), tumor weights were significantly lower than those in the sgNTC group (**Fig. 5J, L; Supplementary Fig. S6A)**. In addition, a weekly decrease in bioluminescence imaging revealed a significant reduction in signal intensity of the sgTAOK3 group compared to the sgNTC control group (**Supplementary Fig. S6B). Supplementary Fig. S6C s**hows loss of *TAOK3* protein expression in sgTAOK3 reproductive tracts compared to those from sgNTC-injected mice.

### *TAOK3* knockdown reshapes the human tumor compartment toward loss of a KRT14-positive invasive program and epithelial differentiation changes

**T**o further understand the function of *TAOK3* in the context of the TME, we employed species-specific RNAseq (SSRS), which uses probabilistic mapping of RNAseq reads to a joint human and mouse transcriptome to assess differential expression in cells of the malignant tumor (human) and the nonmalignant TME (mouse; **Figs. 6 & 7 and Supplementary Fig. S7)**. For each sample, 1.7 × 10^8^ ± 4.3 × 10^7^ reads were generated (**Supplementary Table 4)**, and the average percentage of reads mapping to human (84.5% in sgNTC and 90.1% in sgTAOK3; *p* =0.32; **Supplementary Fig. S7A)** and mouse (15.5% in sgNTC and 9.9% in sgTAOK3; *p* =0.32; **Supplementary Fig. S7B)** did not significantly differ between groups. Differential expression analysis (**Fig. 6A)** showed a balanced set of up- and downregulated genes, with a prominent decline in *KRT14* among the most significantly reduced transcripts (L2FC = −3.42, *p* = 8.82E-22). Pathway enrichment of human DEGs (**Fig. 6B)** highlighted processes tied to epithelial and skin development, regulation of epithelial cell proliferation, extracellular matrix organization, keratinocyte differentiation, and positive regulation of cell migration and MAPK signaling. Consistent with these pathways, immunoblotting (**Fig. 6C)** and immunohistochemistry (**Fig. 6D-F)** confirmed decreased KRT14 protein in sgTAOK3 tumors relative to controls, and multiplex IF in cultured cells demonstrated reduced co-localization of TAOK3 with KRT14 after CRISPRi-mediated *TAOK3* knockdown (**Fig. 6G)**. Of note, KRT14 is expressed in ovarian cancer leader cells, playing a crucial role in ovarian cancer invasion and metastasis while promoting immunosuppression (35,36). Together, these data indicate that epithelial *TAOK3* supports a KRT14-high, invasion-competent differentiation state in the human tumor compartment; its loss downshifts leader cell– associated programs and remodels matrix and motility pathways that underpin local progression.

**Figure 6.**
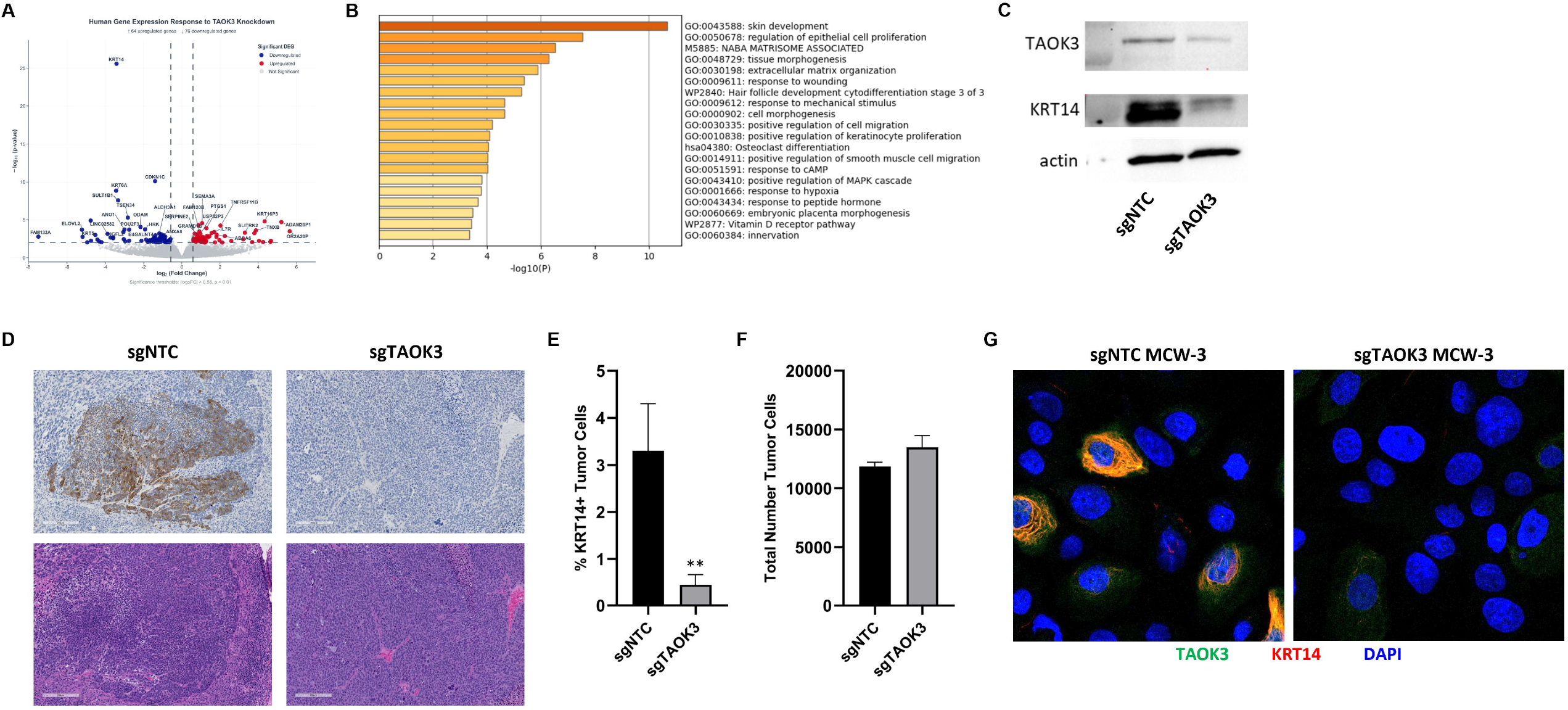
CRISPRi-mediated TAOK3 knockdown reshapes the human tumor compartment toward loss of a KRT14-positive invasive program and epithelial differentiation. **A**, Volcano plot of all significantly downregulated (blue; n =76) and upregulated (red; *n* =64) genes from the human tumor compartment. **B**, Metascape enrichment of human DEGs. **C**, Example western blot images from sgNTC and sgTAOK3-injected mouse reproductive tracts showing decreased TAOK3 and KRT14 protein expression. **D**, Example KRT14 IHC (top panels) and H&E staining (bottom panels) showing increased tumor KRT14 expression in sgNTC vs. sgTAOK3 mice. **E**, Quantification of KRT14+ tumor cells by QuPath confirmed significantly decreased KRT14 expression in sgTAOK3 tumors compared to sgNTC tumors (n=5 for each group). **F**, Total number of cells did not differ across groups. **G**, lmmunocytochemistry confirmed co-staining of *TAOK3* and KRT14 in sgNTC MCW-3 cells which was lost in sgTAOK3 MCW-3 cells. \*\**p*<0.01

### *TAOK3* knockdown remodels the mouse tumor microenvironment toward adaptive immune activation and loss of M2 macrophage polarization

Differentially expressed mouse-compartment (host) transcripts (**Fig. 7A)** from species-specific RNAseq revealed robust immune activation in tumors bearing *TAOK3*-deficient epithelial cells, with enrichment for adaptive immune response, complement/coagulation cascades, leukocyte activation, positive T-cell selection, and phagocytosis pathways (**Fig. 7B)**. A recent report suggested a role for *TAOK3*-JNK signaling in macrophage polarization (37), so we investigated the differential expression of macrophage polarization markers from the mouse compartment. We observed suppression of M2-associated signatures with concordant elevation of M1 programs in sgTAOK3 tumors (**Fig. 7C)**. Quantification of immunofluorescence (Supplementary Fig. S7C-D) of mouse reproductive tracts corroborated a significant reduction of CD206+ (M2) macrophages (**Fig. 7D, E)** without a change in total F4/80+ macrophage number (**Fig. 7F)**, indicating phenotype shift rather than depletion. Finally, to test whether epithelial *TAOK3* directly skews macrophage state, we profiled THP-1 macrophage morphology in co-cultures with sgNTC or sgTAOK3 cervical cancer cells using Incucyte live imaging analysis. Relative to sgNTC co-culture, macrophages exposed to sg*TAOK3* cells displayed significantly higher mean eccentricity (less round/amoeboid—consistent with M1-like activation; **Fig. 7G)** and smaller mean cell size (**Fig. 7H)**, morphometrics that oppose the rounded, enlarged morphology typical of M2 polarization. Together, these data converge on the conclusion that epithelial *TAOK3* promotes an M2-skewed, immunosuppressive niche, showing that loss of *TAOK3* repolarizes macrophages toward an M1-like, phagocytosis-competent phenotype while broadly enhancing adaptive immune pathways.

**Figure 7.**
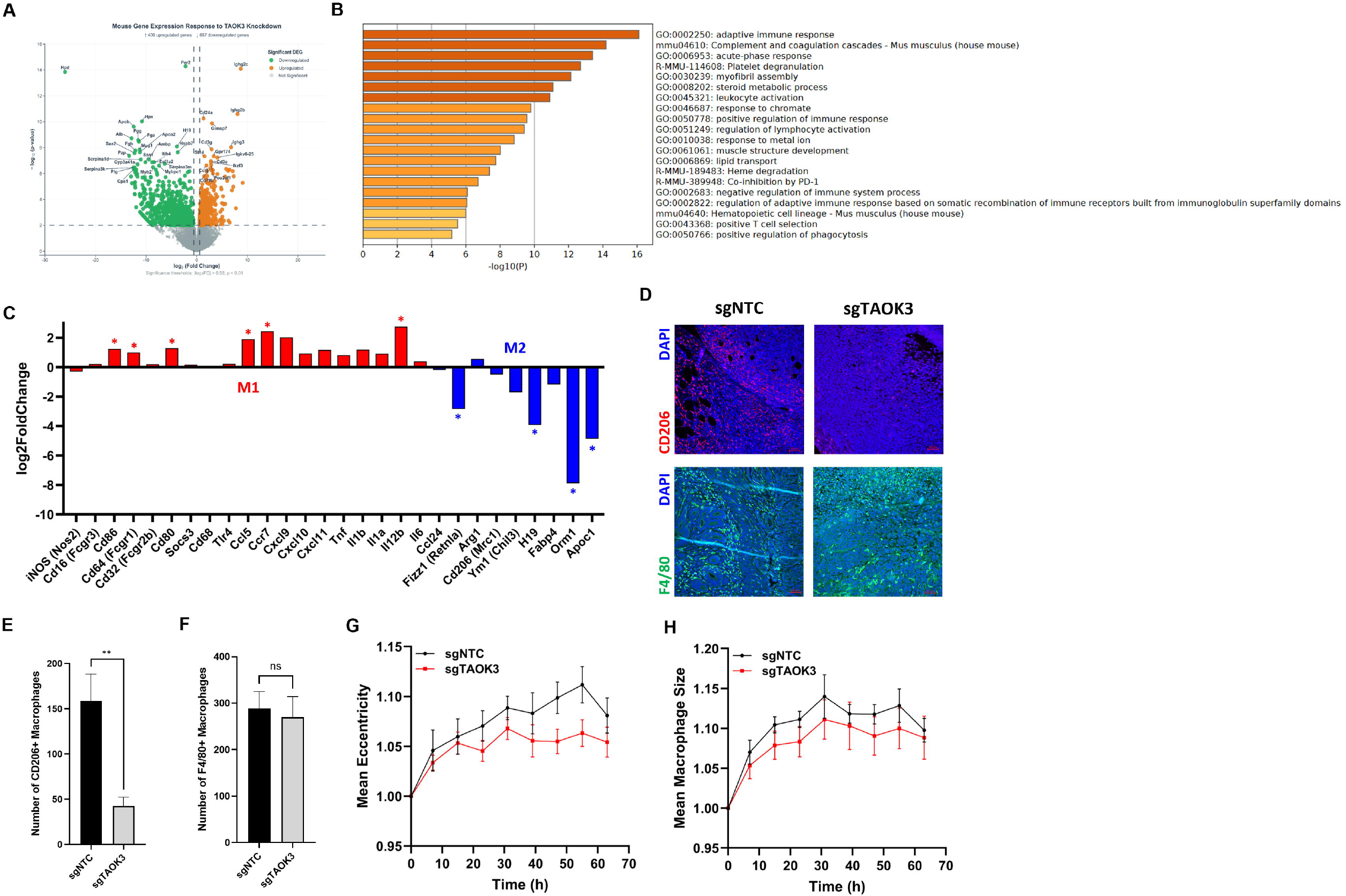
CRISPRi-mediated *TAOK3* knockdown remodels the mouse TME toward adaptive immune activation and loss of M2 macrophage polarization. **A**, Volcano plot of all significantly downregulated (green; n=687) and upregulated (orange; n=400) genes from the mouse tumor microenvironment compartment. **B**, Metascape enrichment of mouse DEGs. **C**, Plot depicting the log2FC of marker genes for Ml (red) and M2 (blue) macrophages from mouse SSRS data shows increase in Ml markers and diminished M2 markers. **D**, Example images from IF of mouse tumors depicting decreased CD206+ (red; M2) macrophages in sgTAOK3 vs. sgNTC mice with no change in total F4/80+ cells (green). QuPath quantification of **E**, number of CD206+ {M2) macrophages and **F**, total F4/80+ cells. Graphing of lncucyte data from CRISPRi SiHa and THP-1 cocultures showing **G**, decreased mean eccentricity and **H**, mean macrophage size of THP-1 cells cocultured with sgTAOK3 vs. sgNTC SiHa. *\*p<0.05; **p<0.01*

### *TAOK3* inhibition triggers methuosis-like vacuolar death with JNK pathway activation and extracellular ATP release

Sustained *TAOK3* suppression induced a pronounced vacuolar phenotype consistent with methuosis-like cell death. In SiHa and MCW-3, time-course imaging with lucifer yellow (fluid-phase endocytosis) and LysoTracker Red (acidic/lysosomal compartments) showed accumulation of large, phase-bright vacuoles that took up lucifer yellow. Across 24–72 hours, vacuoles expanded and remained largely non-acidified/unfused, evidenced by limited colocalization with LysoTracker despite intense lucifer yellow filling (**Fig. 8A-D and Supplementary Fig. S7G)** — hallmarks of defective macropinosome maturation characteristic of methuosis. MOMIPP is a PIKfyve inhibitor shown to induce methuosis and was used as a positive control (38). Further, immunoblotting revealed increased phospho-JNK upon *TAOK3* suppression in both SiHa (**Fig. 8E)** and MCW-3 (**Fig. 8F)**, linking loss of *TAOK3* to activation of stress-activated MAPK signaling, which has been previously linked to methuosis (39). Finally, we surmised that methuotic cell death in *TAOK3*-silenced cells could lead to the release of DAMPs (Damage-Associated Molecular Patterns), which could alter macrophage polarity. Indeed, *TAOK3* knockdown significantly increased extracellular ATP in SiHa (**Fig. 8G)** and MCW-3 (**Fig. 8H)**, consistent with the release of damage-associated signals during vacuolar cell death. Collectively, these findings indicate that *TAOK3* restrains a JNK-linked methuosis, thus sustained *TAOK3* loss or inhibition unleashes vacuolar cytotoxicity and ATP release with potential immunogenic consequences.

**Figure 8.**
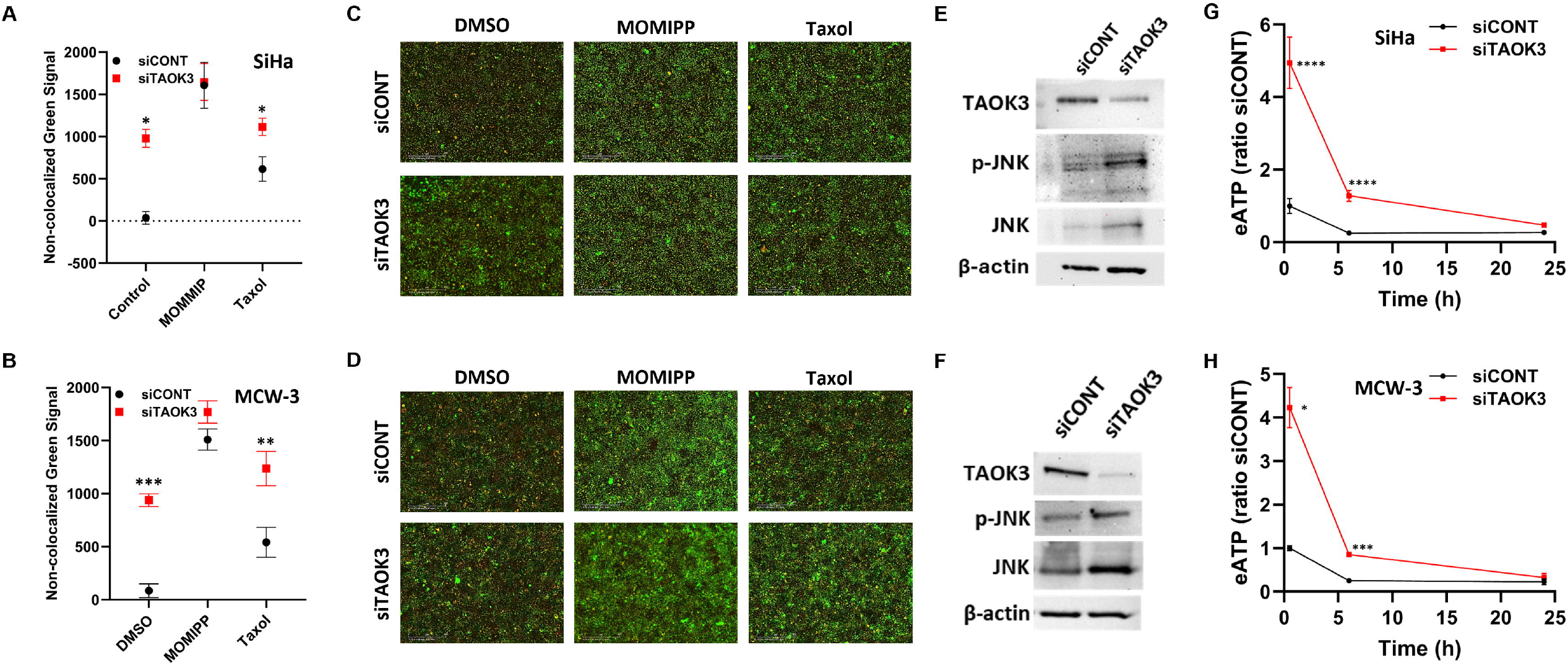
TAOK3 inhibition triggers methuosis-like vacuolar death with JNK pathway activation and extracellular ATP release. **A,C** lncucyte live image quantification of SiHa and MCW-3 **(B, D)** cells shows TAOK3 KD significantly increases the lucifer-yellow positive vacuoles not overlapping with the LysoTracker Red signal which suggests increased methuosis-like cell death in these cells compared to siCONT. MOM IPP (10 µM) used as positive control for methuosis. Treatment with low dose taxol (1 nM) also resulted in significantly more non-lysosome fused vacuoles in siTAOK3 vs. siCONT cells. Consistent with methuosis, JNK pathway activation was increased upon TAOK3 knockdown in SiHa **(E)** and MCW-3 **(F)** cells. Extracellular ATP levels were also significantly increased in TAOK3-silenced SiHa **(G)** and MCW-3 **(H)** cells compared to their control-transfected counterparts. \**p*<0.05; \*\**p*<0.01; \*\*\**p*<0.001; \*\*\*\**p*<0.0001.

## Discussion

This study identifies *TAOK3* as a previously uncharacterized driver of invasive cervical cancer biology. *TAOK3* mRNA and protein were elevated in primary and metastatic ICC relative to normal cervix and localized to a distinct subpopulation of keratin-positive tumor epithelial cells (T3epi). Multi-omic profiling after *TAOK3* silencing showed convergent changes across the transcriptome, proteome, and phosphoproteome, implicating *TAOK3* in cell-cycle progression, cytoskeletal and membrane dynamics, vesicle/endocytic pathways, innate immune signaling, and epithelial differentiation programs. Functionally, *TAOK3* loss or inhibition prolonged G2/M, reduced invasion, and enhanced sensitivity to low-dose paclitaxel *in vitro*. *in vivo, TAOK3* knockdown diminished tumor growth and was associated with an immunopermissive microenvironment characterized by reduced CD206+ M2 macrophages and downregulation of *KRT14*, a marker linked to invasive leader cell states. Together, these data nominate *TAOK3* as a therapeutic target to constrain malignant progression and potentiate chemotherapy in ICC.

TAOK family kinases have been linked to stress-activated MAPK signaling and cytoskeletal regulation in other malignancies (8-10), but their role in cervical cancer was undefined. Our findings place *TAOK3* at a nexus connecting MAPK and Rho GTPase circuitry with endocytosis/vesicle trafficking and actin–membrane remodeling, processes central to epithelial plasticity, EMT, invasion, and metastasis. The T3epi signature identified in scRNA-seq datasets was enriched for cadherin/S100 binding, plasma membrane, vesicles, and leading-edge terms, aligning with a motile, protrusive phenotype. Consistent with this, *TAOK3* knockdown suppressed transcriptional programs related to WNT signaling, mitotic cell cycle, and motility, while upregulating pathways involved in endocytosis, Rho/Miro/RHOBTB3 signaling, and Hippo pathway dysregulation. These changes are congruent with the functional phenotypes of impaired invasion, G2/M delay, and altered vesicle/endocytic trafficking.

An interesting and previously unreported component of the *TAOK3* network is its association with mitochondrial aspartyl-tRNA synthetase 2 *(DARS2). DARS2* mRNA was significantly downregulated in all four *TAOK3*-silenced lines, and DARS2 protein decreased in *TAOK3* KD MCW⍰2 and MCW⍰3 cells. DARS2 is essential for mitochondrial translation and electron transport chain integrity; its loss in hematopoietic stem cells activates a mitochondrial stress response and perturbs iron–sulfur cluster⍰containing respiratory complexes, ultimately compromising proliferative fitness (40). DARS2 upregulation has also been linked to tumorigenesis and poor outcome in several solid tumors (41,42). These observations, together with our data, support a model in which *TAOK3* helps maintain mitochondrial translational capacity and bioenergetic resilience in invasive epithelial cells. Downregulation of *DARS2* after *TAOK3* silencing may therefore contribute to mitochondrial dysfunction, reduced oxidative capacity, and heightened cellular stress in T3epi cells, reinforcing the anti⍰proliferative and pro⍰death effects of *TAOK3* loss.

We also observed consistent downregulation of *PRKCH* across all *TAOK3*-silenced lines. *PRKCH* encodes protein kinase C eta (PKCη), an epithelial⍰enriched PKC isoform implicated in the regulation of adhesion, differentiation, survival, and mechanoresponsive signaling. PKCη promotes invasion and treatment resistance in several epithelial cancers, in part by modulating cytoskeletal organization and Hippo–YAP activity (43). The coordinated suppression of *PRKCH* suggests that *TAOK3* may support an invasion⍰competent, therapy⍰resistant state, at least in part by PKCη⍰dependent control of cortical actin, cell–cell junctions, and transcriptional programs downstream of cytoskeletal cues. Whether PKCη is a direct substrate or transcriptional target of *TAOK3* or lies in a parallel pathway converging on shared cytoskeletal networks will be an important question for future work.

Our *in vivo* data highlight how *TAOK3* orchestrates tumor–microenvironment interactions. In the human tumor compartment, species-specific RNA-seq revealed marked downregulation of *KRT14* and enrichment for pathways involved in epithelial and skin development, keratinocyte differentiation, extracellular matrix organization, and positive regulation of migration and MAPK signaling. *KRT14* marks leader cell populations in ovarian and other cancers (35,36), where it supports collective invasion and immunosuppression. The reduction of *KRT14* protein by immunoblotting and IHC after *TAOK3* knockdown, therefore, suggests loss of an invasion-competent, immunosuppressive epithelial state in ICC. In the mouse compartment, we observed broad enrichment for adaptive immune activation, complement and coagulation cascades, leukocyte activation, T-cell selection, and phagocytosis. Mouse compartment RNAseq and immunofluorescence showed a specific reduction in CD206+ M2 macrophages without loss of total F4/80+ macrophages, indicating repolarization rather than depletion. Co-culture experiments further showed that macrophages exposed to *TAOK3*-deficient tumor cells adopted a more rounded, M1-like morphology with reduced cell size, consistent with transcriptomic evidence of enhanced phagocytic and antigen-presentation programs. These findings fit with emerging evidence that epithelial signaling modules can shape macrophage phenotype via cytokines, chemokines, extracellular vesicles, and DAMPs. In our system, *TAOK3* knockdown reduced expression of several pro-tumoral cytokines and chemokines and altered endocytic and exocytic pathways, any of which could modulate macrophage programming. Additionally, we observed increased HLA class I expression and reductions in antiviral effectors such as RIG-I/DDX58 in some contexts, changes that may rebalance tumor–host interactions away from immune evasion and toward immune surveillance.

Our data also refine the mechanisms by which *TAOK3* intersects with stress⍰activated MAPK signaling and non⍰apoptotic cell death. Prior work has shown that *TAOK3* can activate p38 and/or ERK (8,9) and either inhibit or promote JNK activation, depending on the stimulus and cell type (10), or under endoplasmic reticulum (ER) stress (11). In high⍰*TAOK3* ICC cells, however, prolonged genetic or pharmacologic suppression of *TAOK3* produced a classic methuosis⍰like phenotype: massive lucifer⍰yellow–positive, poorly acidified vacuoles, minimal colocalization with LysoTracker, and progressive loss of viability. This vacuolar death was accompanied by robust JNK phosphorylation and significant increases in extracellular ATP, a canonical damage⍰associated molecular pattern. Methuosis has been linked to dysregulated macropinocytosis and Ras/Rac, Akt–mTOR, and MAPK pathways (44,45). Our observations are consistent with a model in which *TAOK3* normally acts as a “brake”, limiting excessive MAPK/JNK activation and catastrophic macropinocytic vacuolization in tumor cells exposed to high concentrations of growth factors and nutrients, for example, at invasive fronts or perivascular regions. When *TAOK3* is intact, T3epi cells can exploit heightened endocytosis and MAPK signaling to support proliferation, invasion, and survival. When *TAOK3* is lost or inhibited, unchecked stress⍰activated MAPK signaling and endocytic flux may push cells past a threshold into JNK⍰linked methuosis, with release of ATP and other DAMPs that can recruit and activate innate and adaptive immune cells.

The proteomic and phosphoproteomic data further support this view. Enrichment for Rho/Miro/RHOBTB3 signaling, endocytosis, late endosomal microautophagy, and leading-edge terms, together with reduced activity of kinases regulating cytoskeletal projections and ERBB2–ERBB3 signaling, indicates that *TAOK3* coordinates cortical actin, membrane dynamics, and receptor trafficking. *TAOK3* loss altered the phosphorylation of chromatin remodeling and mRNA-processing factors and prolonged G2/M, consistent with roles in checkpoint control and mitotic progression. Upregulation of DNA repair factors such as FAN1 may represent compensatory responses to replication or mitotic stress triggered by *TAOK3* depletion. These alterations create a landscape in which T3epi cells are simultaneously less capable of invasion and mitotic fidelity, yet more prone to MAPK-driven vacuolar death, particularly under conditions of high growth-factor and nutrient flux.

Several limitations should be acknowledged. First, our *in vivo* studies were conducted in nude mice, which lack a fully competent adaptive immune system; the magnitude of *TAOK3*-dependent effects on T-cell–mediated immunity may therefore be underestimated. Second, while our methuosis assays and JNK activation data support a JNK-linked vacuolar death pathway, definitive proof that JNK is required for *TAOK3*-dependent methuosis will require pharmacologic and genetic epistasis experiments. Third, although *DARS2* and PKCη emerged as robust components of the *TAOK3* network, we have not yet tested whether re-expression of either factor rescues specific phenotypes of *TAOK3* loss. Future studies are needed to identify the precise identity of T3+epi cells and their direct downstream *TAOK3* substrates within cytoskeletal, endocytic, and mitotic modules in ICC. Also, further dissection of signaling cascade from *TAOK3* loss to JNK activation and methuosis for identification of ideal time windows for exploiting TAOK3 KD-induced G2M arrest for boosting efficacy of chemotherapy.

In summary, *TAOK3* marks and sustains an invasion-competent epithelial subpopulation in ICC and orchestrates pathways spanning cell-cycle progression, cytoskeleton–membrane dynamics, endocytosis, mitochondrial homeostasis, and tumor–immune interactions. Genetic or pharmacologic *TAOK3* inhibition attenuates proliferation and invasion, sensitizes cells to paclitaxel, and remodels the tumor microenvironment toward reduced M2 macrophage presence and enhanced immune activation, coincident with loss of a *KRT14*-positive leader cell program and the emergence of JNK-linked methuosis. These findings support *TAOK3* as a promising therapeutic target in ICC with the potential to improve chemotherapy efficacy and engage antitumor immunity.

## Author’s Contributions

**M. Iden: C**onceptualization, formal analysis, validation, investigation, visualization, methodology, writing-original draft, and writing-review and editing. **R. Schmidt: I**nvestigation, methodology, and writing-review and editing. **R.D.A.S. Mohammed: F**ormal analysis, investigation, visualization, methodology, writing-original draft, and writing-review and editing. **T.A. Dlugi: V**alidation, investigation, visualization, methodology, writing-original draft, and writing-review and editing. **R. Kumar: D**ata curation, software, formal analysis, visualization, writing-original draft, and writing-review and editing. **S. Tsaih: D**ata curation, software, formal analysis, methodology, writing-review and editing. **B. Nosirov: D**ata curation, software, formal analysis, methodology, writing-original draft, and writing-review and editing. **I.P. Kadamberi: I**nvestigation and visualization. **S. Mittal: I**nvestigation and visualization. **S.L. Narayan: R**esources, data curation, and writing-review and editing. **W.H. Bradley: R**esources and writing-review and editing. **B. Erickson: R**esources and writing-review and editing. **R.C. Czaja: S**upervision and writing-review and editing. **J.C. Felix: S**upervision and writing-review and editing. **V. Jin: M**ethodology and writing-review and editing. **A.I. Ojesina: R**esources, methodology, and writing-review and editing. **S. Pradeep: C**onceptualization, investigation, methodology, writing-original draft, and writing-review and editing. **B.C. Smith: C**onceptualization, methodology, and writing-review and editing. **J.S. Rader: C**onceptualization, resources, supervision, funding acquisition, methodology, project administration, and writing-review and editing.

## Acknowledgements

**W**e would like to express our appreciation for the Children’s Research Institute – Histology Core (RRID:SCR_028287) and Wisconsin Diagnostic Laboratories for their excellent technical assistance and collaboration for histology and immunohistochemistry. Lentivirus generation and flow cytometry for CRISPRi cell lines was performed at the Versiti Blood Research Institute Shared Resources Core Facility (RRID:SCR_025503). Confocal images were acquired at the MCW Oxford Center for Advanced Microscopy – Electron Microscope Core (OxCAM; RRID:SCR_026315). Proteomics and phosphoproteomics experiments were conducted by the MCW Translational Metabolomics Shared Resource (RRID:SCR_027908). This work was supported by the NCI R01CA262198 (to JSR), NCI R01CA279323 (to AIO), NCI R01CA258433 (to SP), and the Women’s Health Research Program (WHRP) in the Department of Obstetrics and Gynecology, Medical College of Wisconsin.

