## Supplemental Table 1 for "*TAOK3* inhibition constrains invasion, potentiates paclitaxel, and reprograms the tumor microenvironment toward anti-tumor immunity in cervical cancer"

| **Supplementary Table 1. Sequences of siRNAs and dual sgRNAs.** | | |
| --- | --- | --- |
| **siRNA Name** | **Qiagen Name (Product Number)** | **Target Sequence** |
| siCONT | AllStars Negative Control siRNA (1027281) | 5’-CAGGGTATCGACGATTACAAA-3’ |
| siTAOK3 | Hs_TAOK3_4 FlexiTube siRNA (Sl00115038) | 5’-AAGAGCGGATCTCCAAACATA-3’ |

| **dual sgRNA** | **Sequence** |
| --- | --- |
| sgNTC | ATTTTGCCCCTGGTTCTTCCACCTTGTTGGGAGTTAAGGCCTCGTCTAGGTTTCAGAGCGAGACGTGCCTGCAGGATACGTCTCAGAAACATGGTGCGGGGGCATGGCCCCGCGTTTAAGAGCTAAGCTGCCAGTTCATTTCTTAGGG |
| sgTAOK3_1.1 | ATTTTGCCCCTGGTTCTTCCACCTTGTTGGCCGGGAGGGACCGGGACAGGTTTCAGAGCGAGACGTGCCTGCAGGATACGTCTCAGAAACATGGTGGAGAGGAGACCCCGGGAGTTTAAGAGCTAAGCTGCCAGTTCATTTCTTAGGG |
