## Supplemental Table 2 for "*TAOK3* inhibition constrains invasion, potentiates paclitaxel, and reprograms the tumor microenvironment toward anti-tumor immunity in cervical cancer"

| **Supplementary Table 2. Sequences of qPCR primers for validating RNAseq DEGs.** | | |
| --- | --- | --- |
| **qPCR Target Gene** | **Forward** **(5'-3')** | **Reverse** **(5'-3')** |
| TAOK3 | GCCATTTTGGCAGAGCAGT | TCTTGAGCCTCATCTAGCCG |
| TCF3 | ACTCCTACAGTGGGCTAGGG | TCTTCTCCTCCTCCGAGTGG |
| APC | AGGCTGCATGAGAGCACTTGTG | CACACTTCCAACTTCTCGCAACG |
| LRP5 | GGACACCAACATGATCGAGTCG | CGCTCAATGCTGTGCAGATTCC |
| ROR1 | GAGGCAACCAAAACACGTCAGAG | GGCACACTCACCCAATTCTTCC |
| PPP3CA | GCCCTGATGAACCAACAGTTCC | GCAGGTGGTTCTTTGAATCGGTC |
| WNT5A | GCTCGCATCCTCATGAACCT | ACCCACCTTGCGGAAGTCT |
| WNT11 | CTGTGAAGGACTCGGAACTCGT | AGCTGTCGCTTCCGTTGGATGT |
| CCND1 | TCTACACCGACAACTCCATCCG | TCTGGCATTTTGGAGAGGAAGTG |
| CSNK2A1 | GGTGAGGATAGCCAAGGTTCTG | TCACTGTGGACAAAGCGTTCCC |
| SKP2 | GATGTGACTGGTCGGTTGCTGT | GAGTTCGATAGGTCCATGTGCTG |
| TIGAR | CCAAAGCAGCCAGGGAAGAGTG | CCGCTTCTTTCAGGATTAGTTGAC |
| DARS2 | CGAGATGAAGGTTCAAGACCAGA | GCCAGGAATACTGGAGCAAACC |
| CXCL8 | TGTAAACATGACTTCCAAGC | AAAACTGCACCTTCACAC |
| IL1A | TGTATGTGACTGCCCAAGATGAAG | AGAGGAGGTTGGTCTCACTACC |
| CSF3 | TCCAGGAGAAGCTGGTGAGTGA | CGCTATGGAGTTGGCTCAAGCA |
| BIRC3 | CGCTTTAAAACATTCTTTAACTGGC | TCTTATCAAGTACTCACACCTTGGA |
| DDX58 | CTCGGAAAATCCCTGCTTTC | GGCCATGTAGCTCAGGATGT |
| RPS18 | TGTGGTGTTGAGGAAAGCA | CTTCAGTCGCTCCAGGTCTT |
