## Supplemental Table 3 for "*TAOK3* inhibition constrains invasion, potentiates paclitaxel, and reprograms the tumor microenvironment toward anti-tumor immunity in cervical cancer"

| **Supplementary Table 3. Antibodies and other reagents used for IHC, ICC, IF, and western blotting.** | | | |  |
| --- | --- | --- | --- | --- |
| **Protein Target** | **Supplier** | **Product Number** | **RRID** | **Assay Concentration** |
| TAOK3 | Sigma Aldrich | HPA017160 | RRID:AB_1851994 | western (1:1000), IHC (1:50) |
| TAOK3 (JIK) | Invitrogen | 712043 | RRID:AB_2784609 | ICC (1:100) |
| TAOK2 | ThermoFisher Scientific | MA5-20685 | RRID:AB_2576583 | western (1:1000) |
| TAOK1 | ThermoFisher Scientific | 703006 | RRID:AB_2809222 | western (1:1000) |
| Cas9 | Sigma Aldrich | SAB4200701 | RRID:AB_2891217 | western (1:1000) |
| TCF3 | ThermoFisher Scientific | 67140-1-IG | RRID:AB_2882439 | western (1:1000) |
| BIRC3 | BioRad | MCA6173 | NA | western (1:1000) |
| CCND1 | ThermoFisher Scientific | MA5-16356 | RRID:AB_2537875 | western (1:1000) |
| β-actin | Cell Signaling Technology | 3700 | RRID:AB_2242334 | western (1:1000) |
| CD206 | Cell Signaling Technology | 24595 | RRID:AB_2892682 | mouse IF (1:400) |
| F4/80 | Cell Signaling Technology | 70076S | RRID:AB_2799771 | mouse IF (1:100) |
| Cytokeratin, Multi (AE1/AE3) | Leica Biosystems | PA0909-U | RRID:AB_2924990 | IHC (1:100) |
| KRT14 | Abcam | AB7800-1001 | RRID:AB_306091 | western (1:1000); IHC (1:50) |
| α-tubulin | Santa Cruz Biotechnology | sc-5286 | RRID:AB_628411 | western (1:1000) |
| Histone H3 | Cell Signaling Technology | 4499S | RRID:AB_10544537 | western (1:1000) |
| Na/K ATPase | Cell Signaling Technology | 3010 | RRID:AB_2060983 | western (1:1000) |
| AlexaFluor™ Plus 647 Phalloidin | Invitrogen | A30107 | NA | ICC (1:1000) |
| DAPI | ThermoFisher Scientific | 62248 | NA | ICC (1:1000) |
