## Supplemental Table 4 for "*TAOK3* inhibition constrains invasion, potentiates paclitaxel, and reprograms the tumor microenvironment toward anti-tumor immunity in cervical cancer"

| **Supplementary Table 4. Percent distribution of human, mouse, and HPV reads in control and TAOK3 knockdown xenograft samples.** | | | | | |
| --- | --- | --- | --- | --- | --- |
| **SampleID** | **Condition** | **Total_TPM_Percentage** | **Percent_Human** | **Percent_Mouse** | **Percent_HPV** |
| B048 | Control (sgNTC) | 100.01 | 73.42 | 26.57 | 0.02 |
| B049 | Control (sgNTC) | 100 | 95 | 4.96 | 0.04 |
| B050 | Control (sgNTC) | 100 | 82.56 | 17.41 | 0.03 |
| B052 | Control (sgNTC) | 99.99 | 86.98 | 12.98 | 0.03 |
| B054 | Experimental (sgTAOK3) | 100 | 85.23 | 14.74 | 0.03 |
| B056 | Experimental (sgTAOK3) | 100 | 92.63 | 7.32 | 0.05 |
| B057 | Experimental (sgTAOK3) | 100 | 89.15 | 10.82 | 0.03 |
| B058 | Experimental (sgTAOK3) | 100 | 93.31 | 6.67 | 0.02 |
