## Supplemental Table 5 for "*TAOK3* inhibition constrains invasion, potentiates paclitaxel, and reprograms the tumor microenvironment toward anti-tumor immunity in cervical cancer"

| **Supplementary Table 5. Genes co-expressed with *TAOK3* in tumor epithelial cells from Fan *et al.*, 2023 dataset.** | | | | | | | | |
| --- | --- | --- | --- | --- | --- | --- | --- | --- |
| **Gene** | **Methods_of_n** | **Frequency** | **Pearson_r** | **Pearson_rank** | **Spearman_r** | **Spearman_rank** | **Kendall_r** | **Kendall_rank** |
| MUC16 | Methods_of_3 | 3 | 0.23668 | 1 | 0.15061 | 4 | 0.11259 | 5 |
| NIBAN2 | Methods_of_3 | 3 | 0.16168 | 13 | 0.11663 | 11 | 0.10212 | 10 |
| RNF213 | Methods_of_3 | 3 | 0.20506 | 2 | 0.11584 | 15 | 0.0963 | 12 |
| SAT1 | Methods_of_3 | 3 | 0.15524 | 15 | 0.17148 | 3 | 0.1154 | 4 |
| TJP2 | Methods_of_3 | 3 | 0.18827 | 4 | 0.11589 | 14 | 0.11197 | 6 |
| ADAM9 | Methods_of_2 | 2 |  |  | 0.11322 | 19 | 0.09177 | 17 |
| ERO1A | Methods_of_2 | 2 |  |  | 0.13481 | 6 | 0.12005 | 3 |
| EZR | Methods_of_2 | 2 |  |  | 0.11538 | 17 | 0.10273 | 9 |
| FNDC3B | Methods_of_2 | 2 |  |  | 0.11529 | 18 | 0.09386 | 14 |
| GPRC5A | Methods_of_2 | 2 |  |  | 0.11223 | 20 | 0.09125 | 19 |
| IGFBP3 | Methods_of_2 | 2 |  |  | 0.14555 | 5 | 0.10683 | 7 |
| LMO7 | Methods_of_2 | 2 | 0.16051 | 14 | 0.1165 | 12 |  |  |
| MALAT1 | Methods_of_2 | 2 |  |  | 0.19034 | 1 | 0.13616 | 2 |
| MUC4 | Methods_of_2 | 2 |  |  | 0.1163 | 13 | 0.09902 | 11 |
| NEAT1 | Methods_of_2 | 2 |  |  | 0.18699 | 2 | 0.14337 | 1 |
| RND3 | Methods_of_2 | 2 |  |  | 0.12692 | 8 | 0.10513 | 8 |
| VMP1 | Methods_of_2 | 2 |  |  | 0.13368 | 7 | 0.09278 | 16 |
| XIST | Methods_of_2 | 2 | 0.1723 | 7 |  |  | 0.08807 | 20 |
| AFDN | Methods_of_1 | 1 |  |  |  |  | 0.09161 | 18 |
| CPEB4 | Methods_of_1 | 1 | 0.14625 | 20 |  |  |  |  |
| EPS8L1 | Methods_of_1 | 1 | 0.16396 | 12 |  |  |  |  |
| F3 | Methods_of_1 | 1 |  |  | 0.12619 | 9 |  |  |
| GABRE | Methods_of_1 | 1 | 0.17963 | 5 |  |  |  |  |
| ICAM1 | Methods_of_1 | 1 | 0.15194 | 17 |  |  |  |  |
| IGHG4 | Methods_of_1 | 1 | 0.17196 | 8 |  |  |  |  |
| IGKC | Methods_of_1 | 1 | 0.20049 | 3 |  |  |  |  |
| IGLC2 | Methods_of_1 | 1 | 0.15042 | 18 |  |  |  |  |
| LCN2 | Methods_of_1 | 1 |  |  | 0.11952 | 10 |  |  |
| LINC00472 | Methods_of_1 | 1 | 0.1642 | 11 |  |  |  |  |
| MXD1 | Methods_of_1 | 1 |  |  | 0.11583 | 16 |  |  |
| PGGHG | Methods_of_1 | 1 | 0.17086 | 9 |  |  |  |  |
| PHLDB2 | Methods_of_1 | 1 | 0.16669 | 10 |  |  |  |  |
| PLCG2 | Methods_of_1 | 1 | 0.15486 | 16 |  |  |  |  |
| PRSS22 | Methods_of_1 | 1 |  |  |  |  | 0.09459 | 13 |
| SLC12A7 | Methods_of_1 | 1 | 0.14739 | 19 |  |  |  |  |
| SPNS2 | Methods_of_1 | 1 |  |  |  |  | 0.09371 | 15 |
| ZNF292 | Methods_of_1 | 1 | 0.17666 | 6 |  |  |  |  |
