## Supplemental Table 6 for "*TAOK3* inhibition constrains invasion, potentiates paclitaxel, and reprograms the tumor microenvironment toward anti-tumor immunity in cervical cancer"

| **Supplementary Table 6. Genes co-expressed with *TAOK3* in tumor epithelial cells from Terekhanova *et al.*, 2023 dataset.** | | | | | | | | | | |
| --- | --- | --- | --- | --- | --- | --- | --- | --- | --- | --- |
| **Gene** | **Methods_of_n** | **Frequency** | **Pearson_r** | **Pearson_rank** | **Spearman_r** | **Spearman_rank** | **Kendall_r** | **Kendall_rank** | **CSCORE_r** | **CSCORE_rank** |
| PLAC8 | Methods_of_4 | 4 | 0.16979 | 5 | 0.08626 | 12 | 0.0883 | 12 | 0.40546 | 12 |
| ABLIM3 | Methods_of_3 | 3 | 0.19206 | 1 | 0.08729 | 11 | 0.09816 | 7 |  |  |
| DUOX2 | Methods_of_3 | 3 | 0.13939 | 10 | 0.08901 | 9 | 0.0918 | 11 |  |  |
| PPL | Methods_of_3 | 3 | 0.14799 | 8 | 0.08605 | 13 | 0.08709 | 13 |  |  |
| SCEL | Methods_of_3 | 3 | 0.18249 | 3 | 0.10495 | 6 | 0.11185 | 2 |  |  |
| SPINK5 | Methods_of_3 | 3 | 0.12986 | 12 | 0.11068 | 4 | 0.12874 | 1 |  |  |
| SPNS2 | Methods_of_3 | 3 | 0.17259 | 4 | 0.08828 | 10 | 0.08342 | 16 |  |  |
| SPRR3 | Methods_of_3 | 3 | 0.19119 | 2 | 0.11106 | 3 | 0.10743 | 3 |  |  |
| TPRG1 | Methods_of_3 | 3 | 0.12829 | 16 | 0.08224 | 14 | 0.08675 | 14 |  |  |
| AHNAK | Methods_of_2 | 2 |  |  | 0.10455 | 7 | 0.10664 | 4 |  |  |
| CSMD1 | Methods_of_2 | 2 |  |  | 0.11764 | 2 | 0.09745 | 8 |  |  |
| CSTA | Methods_of_2 | 2 |  |  | 0.10032 | 8 | 0.08269 | 19 |  |  |
| MUC20 | Methods_of_2 | 2 |  |  | 0.0745 | 20 | 0.1 | 6 |  |  |
| PLEKHM1 | Methods_of_2 | 2 | 0.124 | 20 |  |  |  |  | 0.42357 | 8 |
| PRRC2C | Methods_of_2 | 2 |  |  | 0.12324 | 1 | 0.10296 | 5 |  |  |
| RHBDL2 | Methods_of_2 | 2 |  |  | 0.0749 | 19 | 0.08201 | 20 |  |  |
| RNF213 | Methods_of_2 | 2 |  |  | 0.10849 | 5 | 0.08305 | 18 |  |  |
| TMPRSS11E | Methods_of_2 | 2 | 0.15122 | 7 |  |  | 0.08308 | 17 |  |  |
| TMPRSS2 | Methods_of_2 | 2 | 0.15189 | 6 |  |  |  |  | 0.39922 | 14 |
| UBE2H | Methods_of_2 | 2 | 0.12916 | 14 |  |  |  |  | 0.40894 | 10 |
| ADGRF1 | Methods_of_1 | 1 |  |  |  |  | 0.09379 | 9 |  |  |
| AHNAK2 | Methods_of_1 | 1 |  |  |  |  | 0.09275 | 10 |  |  |
| ANKRD13A | Methods_of_1 | 1 |  |  |  |  |  |  | 0.43691 | 7 |
| ANXA11 | Methods_of_1 | 1 |  |  |  |  |  |  | 0.41226 | 9 |
| APOBEC3A | Methods_of_1 | 1 | 0.13418 | 11 |  |  |  |  |  |  |
| ARHGAP27 | Methods_of_1 | 1 |  |  |  |  |  |  | 0.47532 | 3 |
| BCAT1 | Methods_of_1 | 1 | 0.1243 | 19 |  |  |  |  |  |  |
| BCL2L1 | Methods_of_1 | 1 |  |  |  |  |  |  | 0.45266 | 5 |
| CASTOR2 | Methods_of_1 | 1 |  |  |  |  |  |  | 0.37556 | 19 |
| CDKL5 | Methods_of_1 | 1 |  |  |  |  |  |  | 0.48803 | 2 |
| CYP3A5 | Methods_of_1 | 1 | 0.12593 | 18 |  |  |  |  |  |  |
| DHRS9 | Methods_of_1 | 1 | 0.12768 | 17 |  |  |  |  |  |  |
| DNAJC5 | Methods_of_1 | 1 |  |  | 0.07521 | 18 |  |  |  |  |
| EPS8L1 | Methods_of_1 | 1 | 0.1293 | 13 |  |  |  |  |  |  |
| LNX1 | Methods_of_1 | 1 |  |  |  |  |  |  | 0.40798 | 11 |
| NXN | Methods_of_1 | 1 |  |  |  |  | 0.084 | 15 |  |  |
| PADI1 | Methods_of_1 | 1 |  |  | 0.08036 | 15 |  |  |  |  |
| PICALM | Methods_of_1 | 1 |  |  |  |  |  |  | 0.44219 | 6 |
| RMND5A | Methods_of_1 | 1 |  |  |  |  |  |  | 0.46371 | 4 |
| RNF169 | Methods_of_1 | 1 |  |  |  |  |  |  | 0.38946 | 16 |
| SERPINB13 | Methods_of_1 | 1 |  |  | 0.07907 | 16 |  |  |  |  |
| SLC12A6 | Methods_of_1 | 1 |  |  |  |  |  |  | 0.38835 | 17 |
| SORT1 | Methods_of_1 | 1 |  |  |  |  |  |  | 0.50446 | 1 |
| SPECC1 | Methods_of_1 | 1 |  |  |  |  |  |  | 0.3665 | 20 |
| SPRR2A | Methods_of_1 | 1 | 0.14303 | 9 |  |  |  |  |  |  |
| STS | Methods_of_1 | 1 |  |  |  |  |  |  | 0.40186 | 13 |
| SULT2B1 | Methods_of_1 | 1 |  |  | 0.07621 | 17 |  |  |  |  |
| TMPRSS11B | Methods_of_1 | 1 | 0.12854 | 15 |  |  |  |  |  |  |
| USP6NL | Methods_of_1 | 1 |  |  |  |  |  |  | 0.38347 | 18 |
| ZSWIM4 | Methods_of_1 | 1 |  |  |  |  |  |  | 0.3905 | 15 |
