## Supplemental Table 7 for "*TAOK3* inhibition constrains invasion, potentiates paclitaxel, and reprograms the tumor microenvironment toward anti-tumor immunity in cervical cancer"

| source | term_name | term_id | adjusted_p_value | negative_log10_of_adjusted_p_value | term_size | query_size | intersection_size | effective_domain_size | intersections |
| --- | --- | --- | --- | --- | --- | --- | --- | --- | --- |
| GO:MF | cadherin binding | GO:0045296 | 0.000434 | 3.3623265 | 335 | 80 | 10 | 20208 | NIBAN2,TJP2,EZR,GPRC5A,AFDN,EPS8L1,PHLDB2,PPL,AHNAK,PICALM |
| GO:MF | cell adhesion molecule binding | GO:0050839 | 0.001123 | 2.9497908 | 563 | 80 | 12 | 20208 | NIBAN2,TJP2,ADAM9,EZR,GPRC5A,ICAM1,AFDN,EPS8L1,PHLDB2,PPL,AHNAK,PICALM |
| GO:MF | S100 protein binding | GO:0044548 | 0.01133 | 1.9457689 | 14 | 80 | 3 | 20208 | EZR,AHNAK,ANXA11 |
| GO:MF | serine-type endopeptidase activity | GO:0004252 | 0.048541 | 1.3138912 | 184 | 80 | 6 | 20208 | F3,PRSS22,RHBDL2,TMPRSS11E,TMPRSS2,TMPRSS11B |
| GO:BP | epithelial cell differentiation | GO:0030855 | 0.038092 | 1.4191633 | 745 | 79 | 12 | 20972 | TJP2,ADAM9,EZR,ICAM1,AFDN,PPL,SCEL,SPINK5,SPRR3,CSTA,DHRS9,SULT2B1 |
| GO:CC | plasma membrane | GO:0005886 | 1.56E-05 | 4.8072851 | 5906 | 82 | 45 | 22155 | MUC16,NIBAN2,TJP2,ADAM9,EZR,GPRC5A,LMO7,MUC4,RND3,VMP1,F3,ICAM1,AFDN,EPS8L1,GABRE,IGHG4,IGKC,IGLC2,PHLDB2,PLCG2,SLC12A7,SPNS2,DUOX2,PPL,SCEL,SPRR3,AHNAK,CSTA,MUC20,RHBDL2,TMPRSS11E,TMPRSS2,ADGRF1,AHNAK2,ANKRD13A,ANXA11,ARHGAP27,CDKL5,DNAJC5,PICALM,SLC12A6,SORT1,SPRR2A,TMPRSS11B,USP6NL |
| GO:CC | cell periphery | GO:0071944 | 3.1E-05 | 4.5090334 | 6263 | 82 | 46 | 22155 | MUC16,NIBAN2,TJP2,ADAM9,EZR,GPRC5A,LMO7,MUC4,RND3,VMP1,F3,ICAM1,AFDN,EPS8L1,GABRE,IGHG4,IGKC,IGLC2,PHLDB2,PLCG2,SLC12A7,SPNS2,DUOX2,PPL,SCEL,SPINK5,SPRR3,AHNAK,CSTA,MUC20,RHBDL2,TMPRSS11E,TMPRSS2,ADGRF1,AHNAK2,ANKRD13A,ANXA11,ARHGAP27,CDKL5,DNAJC5,PICALM,SLC12A6,SORT1,SPRR2A,TMPRSS11B,USP6NL |
| GO:CC | membrane | GO:0016020 | 0.000187 | 3.7291761 | 9975 | 82 | 59 | 22155 | MUC16,NIBAN2,RNF213,TJP2,ADAM9,FNDC3B,ERO1A,EZR,GPRC5A,LMO7,MUC4,RND3,VMP1,F3,ICAM1,AFDN,EPS8L1,GABRE,IGHG4,IGKC,IGLC2,PHLDB2,PLCG2,PRSS22,SLC12A7,SPNS2,DUOX2,PPL,SCEL,SPINK5,SPRR3,AHNAK,CSMD1,CSTA,MUC20,PLEKHM1,PRRC2C,RHBDL2,TMPRSS11E,TMPRSS2,ADGRF1,AHNAK2,ANKRD13A,ANXA11,ARHGAP27,BCL2L1,CDKL5,CYP3A5,DHRS9,DNAJC5,PICALM,SERPINB13,SLC12A6,SORT1,SPECC1,SPRR2A,STS,TMPRSS11B,USP6NL |
| GO:CC | vesicle | GO:0031982 | 0.000214 | 3.670363 | 4032 | 82 | 34 | 22155 | MUC16,NIBAN2,ADAM9,EZR,GPRC5A,MUC4,ICAM1,LCN2,EPS8L1,IGHG4,IGLC2,PLCG2,SPNS2,PLAC8,DUOX2,PPL,SCEL,SPINK5,SPRR3,AHNAK,PLEKHM1,TMPRSS2,ADGRF1,ANKRD13A,ANXA11,BCL2L1,DNAJC5,PICALM,SERPINB13,SORT1,SPRR2A,SULT2B1,TMPRSS11B,USP6NL |
| GO:CC | cornified envelope | GO:0001533 | 0.000772 | 3.1121526 | 60 | 82 | 5 | 22155 | PPL,SCEL,SPRR3,CSTA,SPRR2A |
| GO:CC | extracellular exosome | GO:0070062 | 0.001517 | 2.8188797 | 2111 | 82 | 22 | 22155 | MUC16,NIBAN2,ADAM9,EZR,GPRC5A,MUC4,ICAM1,LCN2,EPS8L1,IGHG4,IGLC2,PLCG2,DUOX2,PPL,SCEL,SPRR3,AHNAK,TMPRSS2,ANXA11,SERPINB13,SULT2B1,TMPRSS11B |
| GO:CC | extracellular vesicle | GO:1903561 | 0.001905 | 2.7200147 | 2141 | 82 | 22 | 22155 | MUC16,NIBAN2,ADAM9,EZR,GPRC5A,MUC4,ICAM1,LCN2,EPS8L1,IGHG4,IGLC2,PLCG2,DUOX2,PPL,SCEL,SPRR3,AHNAK,TMPRSS2,ANXA11,SERPINB13,SULT2B1,TMPRSS11B |
| GO:CC | extracellular membrane-bounded organelle | GO:0065010 | 0.00192 | 2.7167528 | 2142 | 82 | 22 | 22155 | MUC16,NIBAN2,ADAM9,EZR,GPRC5A,MUC4,ICAM1,LCN2,EPS8L1,IGHG4,IGLC2,PLCG2,DUOX2,PPL,SCEL,SPRR3,AHNAK,TMPRSS2,ANXA11,SERPINB13,SULT2B1,TMPRSS11B |
| GO:CC | extracellular organelle | GO:0043230 | 0.00192 | 2.7167528 | 2142 | 82 | 22 | 22155 | MUC16,NIBAN2,ADAM9,EZR,GPRC5A,MUC4,ICAM1,LCN2,EPS8L1,IGHG4,IGLC2,PLCG2,DUOX2,PPL,SCEL,SPRR3,AHNAK,TMPRSS2,ANXA11,SERPINB13,SULT2B1,TMPRSS11B |
| GO:CC | extracellular region | GO:0005576 | 0.002217 | 2.6542185 | 4249 | 82 | 33 | 22155 | MUC16,NIBAN2,ADAM9,ERO1A,EZR,GPRC5A,MUC4,IGFBP3,F3,ICAM1,LCN2,EPS8L1,IGHG4,IGLC2,PLCG2,PRSS22,PLAC8,DUOX2,PPL,SCEL,SPINK5,SPRR3,AHNAK,CSTA,MUC20,TMPRSS11E,TMPRSS2,ADGRF1,ANXA11,SERPINB13,SPRR2A,SULT2B1,TMPRSS11B |
| GO:CC | cell junction | GO:0030054 | 0.002859 | 2.5438475 | 2370 | 82 | 23 | 22155 | NIBAN2,TJP2,ADAM9,EZR,LMO7,RND3,ICAM1,AFDN,CPEB4,GABRE,PHLDB2,SLC12A7,ABLIM3,DUOX2,PPL,AHNAK,BCL2L1,CDKL5,DNAJC5,LNX1,PICALM,SLC12A6,SORT1 |
| GO:CC | cytosol | GO:0005829 | 0.002948 | 2.5304943 | 5607 | 82 | 39 | 22155 | NIBAN2,RNF213,TJP2,SAT1,EZR,LMO7,RND3,AFDN,EPS8L1,MXD1,PGGHG,PHLDB2,PLCG2,DUOX2,PPL,SPINK5,SPRR3,AHNAK,CSTA,PRRC2C,UBE2H,AHNAK2,ANXA11,ARHGAP27,BCAT1,BCL2L1,CASTOR2,DNAJC5,NXN,PADI1,PICALM,RMND5A,RNF169,SERPINB13,SORT1,SPECC1,SPRR2A,SULT2B1,USP6NL |
| GO:CC | extracellular space | GO:0005615 | 0.006814 | 2.1666083 | 3252 | 82 | 27 | 22155 | MUC16,NIBAN2,ADAM9,ERO1A,EZR,GPRC5A,MUC4,IGFBP3,F3,ICAM1,LCN2,EPS8L1,IGHG4,IGLC2,PLCG2,PRSS22,DUOX2,PPL,SCEL,SPRR3,AHNAK,CSTA,TMPRSS2,ANXA11,SERPINB13,SULT2B1,TMPRSS11B |
| GO:CC | anchoring junction | GO:0070161 | 0.034361 | 1.4639332 | 911 | 82 | 12 | 22155 | NIBAN2,TJP2,ADAM9,EZR,LMO7,RND3,ICAM1,AFDN,PHLDB2,DUOX2,PPL,AHNAK |
| GO:CC | cell leading edge | GO:0031252 | 0.04637 | 1.3337644 | 423 | 82 | 8 | 22155 | EZR,EPS8L1,GABRE,PHLDB2,PLCG2,ABLIM3,DUOX2,CDKL5 |
| KEGG | Steroid hormone biosynthesis | KEGG:00140 | 0.01707 | 1.7677716 | 66 | 39 | 4 | 8716 | CYP3A5,DHRS9,STS,SULT2B1 |
| REAC | Dectin-2 family | REAC:R-HSA-5621480 | 0.001421 | 2.8473189 | 26 | 46 | 4 | 11056 | MUC16,MUC4,PLCG2,MUC20 |
| REAC | Defective GALNT3 causes HFTC | REAC:R-HSA-5083625 | 0.014087 | 1.8511699 | 16 | 46 | 3 | 11056 | MUC16,MUC4,MUC20 |
| REAC | Defective GALNT12 causes CRCS1 | REAC:R-HSA-5083636 | 0.014087 | 1.8511699 | 16 | 46 | 3 | 11056 | MUC16,MUC4,MUC20 |
| REAC | Defective C1GALT1C1 causes TNPS | REAC:R-HSA-5083632 | 0.017056 | 1.768116 | 17 | 46 | 3 | 11056 | MUC16,MUC4,MUC20 |
| REAC | Termination of O-glycan biosynthesis | REAC:R-HSA-977068 | 0.043651 | 1.3600048 | 23 | 46 | 3 | 11056 | MUC16,MUC4,MUC20 |
| MIRNA | hsa-mir-527 | MIRNA:hsa-mir-527 | 0.000319 | 3.4961417 | 1413 | 73 | 21 | 16638 | RNF213,ADAM9,FNDC3B,EZR,RND3,VMP1,CPEB4,MXD1,PHLDB2,SLC12A7,ZNF292,AHNAK,PLEKHM1,PRRC2C,UBE2H,ANKRD13A,ARHGAP27,BCAT1,DNAJC5,PADI1,RMND5A |
| MIRNA | hsa-mir-518a-5p | MIRNA:hsa-mir-518a-5p | 0.000327 | 3.4859404 | 1415 | 73 | 21 | 16638 | RNF213,ADAM9,FNDC3B,EZR,RND3,VMP1,CPEB4,MXD1,PHLDB2,SLC12A7,ZNF292,AHNAK,PLEKHM1,PRRC2C,UBE2H,ANKRD13A,ARHGAP27,BCAT1,DNAJC5,PADI1,RMND5A |
| MIRNA | hsa-mir-4253 | MIRNA:hsa-mir-4253 | 0.027642 | 1.5584312 | 911 | 73 | 14 | 16638 | RNF213,FNDC3B,GPRC5A,ICAM1,GABRE,PHLDB2,SLC12A7,PLAC8,MUC20,TMPRSS2,UBE2H,BCL2L1,RNF169,SORT1 |
| MIRNA | hsa-mir-98-5p | MIRNA:hsa-mir-98-5p | 0.037027 | 1.4314784 | 698 | 73 | 12 | 16638 | FNDC3B,LMO7,RND3,ICAM1,GABRE,MXD1,PLCG2,PRSS22,SLC12A7,PRRC2C,TMPRSS2,PICALM |
| HPA | Skin 2; cells in granular layer | HPA:0470181 | 0.018916 | 1.7231739 | 719 | 51 | 12 | 11007 | TJP2,SAT1,EZR,MXD1,PPL,SCEL,SPINK5,CSTA,AHNAK2,PICALM,SERPINB13,SULT2B1 |
