## Supplemental Table 8 for "*TAOK3* inhibition constrains invasion, potentiates paclitaxel, and reprograms the tumor microenvironment toward anti-tumor immunity in cervical cancer"

| **Supplementary Table 8. Common DEGs in four unique *TAOK3* KD cervical cancer cell lines.** | | |
| --- | --- | --- |
| **DOWN** |  | **UP** |
| ENSG00000134982 |  | ENSG00000131943 |
| ENSG00000162885 |  | ENSG00000185015 |
| ENSG00000118276 |  | ENSG00000091527 |
| ENSG00000023445 |  | ENSG00000169826 |
| ENSG00000184992 |  | ENSG00000188215 |
| ENSG00000127995 |  | ENSG00000170456 |
| ENSG00000149231 |  | ENSG00000121644 |
| ENSG00000110092 |  | ENSG00000136048 |
| ENSG00000123219 |  | ENSG00000183161 |
| ENSG00000188419 |  | ENSG00000100522 |
| ENSG00000065615 |  | ENSG00000235750 |
| ENSG00000117593 |  | ENSG00000153714 |
| ENSG00000107201 |  | ENSG00000107789 |
| ENSG00000024526 |  | ENSG00000160588 |
| ENSG00000163577 |  | ENSG00000163293 |
| ENSG00000118985 |  | ENSG00000156675 |
| ENSG00000134531 |  | ENSG00000188725 |
| ENSG00000087502 |  | ENSG00000160075 |
| ENSG00000116199 |  | ENSG00000112079 |
| ENSG00000157107 |  | ENSG00000196116 |
| ENSG00000052795 |  | ENSG00000196663 |
| ENSG00000139926 |  |  |
| ENSG00000138757 |  |  |
| ENSG00000115159 |  |  |
| ENSG00000111906 |  |  |
| ENSG00000010404 |  |  |
| ENSG00000136381 |  |  |
| ENSG00000050130 |  |  |
| ENSG00000102781 |  |  |
| ENSG00000174010 |  |  |
| ENSG00000143815 |  |  |
| ENSG00000145685 |  |  |
| ENSG00000163428 |  |  |
| ENSG00000173926 |  |  |
| ENSG00000140943 |  |  |
| ENSG00000106484 |  |  |
| ENSG00000101752 |  |  |
| ENSG00000178802 |  |  |
| ENSG00000120333 |  |  |
| ENSG00000105887 |  |  |
| ENSG00000204899 |  |  |
| ENSG00000124357 |  |  |
| ENSG00000171208 |  |  |
| ENSG00000164164 |  |  |
| ENSG00000131779 |  |  |
| **DOWN (continued)**  ENSG00000138814 |  |  |
| ENSG00000162409 |  |  |
| ENSG00000027075 |  |  |
| ENSG00000171016 |  |  |
| ENSG00000163694 |  |  |
| ENSG00000135249 |  |  |
| ENSG00000185483 |  |  |
| ENSG00000163904 |  |  |
| ENSG00000187231 |  |  |
| ENSG00000164054 |  |  |
| ENSG00000116991 |  |  |
| ENSG00000145604 |  |  |
| ENSG00000143570 |  |  |
| ENSG00000163683 |  |  |
| ENSG00000135587 |  |  |
| ENSG00000021574 |  |  |
| ENSG00000196369 |  |  |
| ENSG00000136854 |  |  |
| ENSG00000198252 |  |  |
| ENSG00000135090 |  |  |
| ENSG00000071564 |  |  |
| ENSG00000163444 |  |  |
| ENSG00000133678 |  |  |
| ENSG00000167904 |  |  |
| ENSG00000120802 |  |  |
| ENSG00000125247 |  |  |
| ENSG00000183864 |  |  |
| ENSG00000165832 |  |  |
| ENSG00000196236 |  |  |
| ENSG00000143324 |  |  |
| ENSG00000180233 |  |  |
